# LrhA promotes CRISPR-Cas immunity by enhancing primed adaptation in *Escherichia coli*

**DOI:** 10.1101/2023.08.14.552655

**Authors:** Mengdie Fang, Na Li, Mengxin Gong, Mingyue Fei, Yalan Lu, Mingjing Yu, Fabai Wu, Dongchang Sun

## Abstract

The CRISPR-based defense system protects prokaryotes against foreign genetic elements. Previous studies indicated that the activity of the CRISPR-Cas system is the outcome of the combined effect of adaptation and interference. However, it remains unclear how coordination of the two processes is regulated to optimize the efficacy of adaptive immunity. Here, we identify the LysR-type transcriptional regulator LrhA as a new CRISPR-Cas activator which plays an important role in coordinating adaptation and interference. Through directly binding to the promoter of the *cas* operon, LrhA enhances the CRISPR-Cas adaptive immunity against bacteriophage infection by stimulating the transcription of *cas* genes. Moderate activation of *cas* genes by LrhA efficiently promoted the clearance of horizontally transferred CRISPR-targeted plasmid via increasing the acquisition of new spacers by interference-driven adaptation in *Escherichia coli*. Our results indicate that intermediate level of transcription of *cas* genes achieved optimal outcomes of the adaptive immunity through triggering positive feedback between adaptation and interference, highlighting the importance of fine-tuning expression of CRISPR-Cas system.

## INTRODUCTION

Prokaryotes have evolved diverse defense systems against plasmids and viruses ^1, 2, 3,4^. Among them, the CRISPR-Cas adaptive immune system provides bacteria and archaea with acquired and inheritable immunity against previously encountered invading nucleic acids^5, 6, 7, 8, 9, 10^. The CRISPR-Cas system acts in three distinct stages. During the first contact with a foreign genetic element, short DNA fragments from this element are incorporated into the CRISPR array, which contains the genetic memory of previously infecting elements^11, 12, 13^. The incorporation of a new spacer derived from the foreign DNA leads to the expansion of the size of the CRISPR array. This process is termed adaptation^14, 15, 16, 17^. Under certain conditions, Cas proteins are expressed, accompanied by the transcription of the CRISPR array, and subsequent processing of the premature CRISPR RNA (pre-crRNA) into mature crRNAs^18, 19^. In the final stage termed interference, crRNAs guide Cas proteins to the target sequence, where the Cas effectors are recruited for cleaving the previously encountered foreign DNA^20, 21^. Native CRISPR-Cas systems have been engineered for broad applications such as genome editing tools^22, 23, 24, 25^ and antimicrobials for combating multi-drug resistant (MDR) bacteria ^26, 27, 28, 29, 30, 31^.

CRISPR-Cas adaptive immunity represents a powerful weapon of defense in prokaryotes. On the other hand, invasive nucleic acids often rapidly evolve resistance to CRISPR immunity by mutating the CRISPR-targeted region. To fight back, a process named ‘primed adaptation’ is provoked to rapidly incorporate new spacers, derived from DNA fragments produced by interference, for upgrading the CRISPR-Cas adaptive immunity^32, 33, 34^. Primed adaptation requires components for assembly of the interference and the adaptation complexes^32, 35^. Nevertheless, a chronic high-level of CRISPR-Cas adaptive immunity increases the burden of producing Cas proteins and crRNAs, as well as the risk of suicide from autoimmunity^36, 37, 38, 39^. To finely tune the CRISPR immunity to a moderate level, interference and adaptation are often inversely correlated: an intermediate level of interference allows invasive nucleic acids to persist longer, leading to higher levels of adaptation (interference-driven adaptation), while a strong interference leads to a relatively low level of adaptation ^40, 41^. Although the working mechanisms of CRISPR adaptation and interference have been intensively investigated^11, 14, 16, 17, 35, 42, 43, 44, 45, 46, 47, 48^, little is known about how the CRISPR-Cas adaptive immunity is adjusted by leveraging both CRISPR interference and adaptation, when facing different types of vulnerabilities ^18, 19^.

The type I CRISPR-Cas system, comprised of the multi-subunit RNA-guided surveillance complex Cascade, the Cas3 nuclease, and the Cas1-Cas2 integrase, is most prevalent in prokaryotes^49^. Regulation studies have focused on type I CRISPR-Cas systems ^18, 40, 50, 51, 52, 53, 54, 55, 56, 57, 58, 59^, especially the type I-E CRISPR-Cas system of *Escherichia coli*, in which CRISPR interference is regulated by the LysR type transcriptional regulator (LTTR) LeuO ^60, 61^, the histone-like proteins H-NS and StpA ^62, 63, 64^, and the carbon catabolite regulator CRP^65^. Nevertheless, regulation of CRISPR adaptation has rarely been investigated in *E. coli*. Less is known about how CRISPR adaptation and interference are coordinated. We hypothesized the presence of host factor(s) that are involved in the coordination of CRISPR adaptation and interference, and effectively aid the bacteria to counter foreign DNA under certain scenarios.

To search for host factor(s) that control the adaptive immunity through coordinating CRISPR adaptation and interference in *E. coli*, we screened for new regulators involved in regulating transcription of *cas* genes by combining DNA pull-down and mass spectrometry, and identified the LTTR protein LrhA as an important activator, which directly bound to the promoter of the *cas* operon, and stimulated the CRISPR-Cas adaptive immunity against bacteriophage infection. By promoting a positive feedback circuit between interference and adaptation, LrhA remarkably augmented the clearance of CRISPR-targeted plasmid, via moderately stimulating the transcription of *cas* genes required for both adaptation and interference. Our work uncovers that intermediate level of transcription of *cas* genes enables coordination of interference and adaptation to achieve optimal adaptive immunity, gaining new insights into the regulation of bacterial defence against horizontally transferred plasmids.

## RESULTS

### Screening for proteins binding to the promoter of *cas* genes

Expression of the *cas* operon is regulated by H-NS and LeuO, which bind to specific sites within P*_cas_* ^60, 62^ (Fig. 1a, Fig. S1). To screen for new regulators that potentially affect the transcription of *cas* genes, we performed a DNA pull-down assay using a DNA probe containing P*_cas_* as a bait (Fig. 1a). Because H-NS binds to high-affinity sites within P*_cas_* and possibly spreads along the adjacent DNA^66^, potentially masking the binding sites of other regulators, we screened for proteins binding to P*_cas_* in *hns*-deletion mutants of two *E. coli* strains (MC4100 and BW25113). Mass spectrometry analysis showed that more than one hundred proteins in each strain were isolated with the biotinylated P*_cas_* (Fig. S2a). Among them, 49 proteins were detected in both strains (Fig. S2b, Table S1). According to the bioinformatic analysis of proteins with high coverage of matched peptides, 18 proteins were classified as transcriptional factors (TFs) (Fig. 1b, Table S2). Two previously documented regulators of the CRISPR-Cas system (i.e. StpA^63, 64^ and SlyA^67^) were identified in the protein mixture bound to P*_cas_,* indicating that the screening approach was effective.

**Figure 1.**
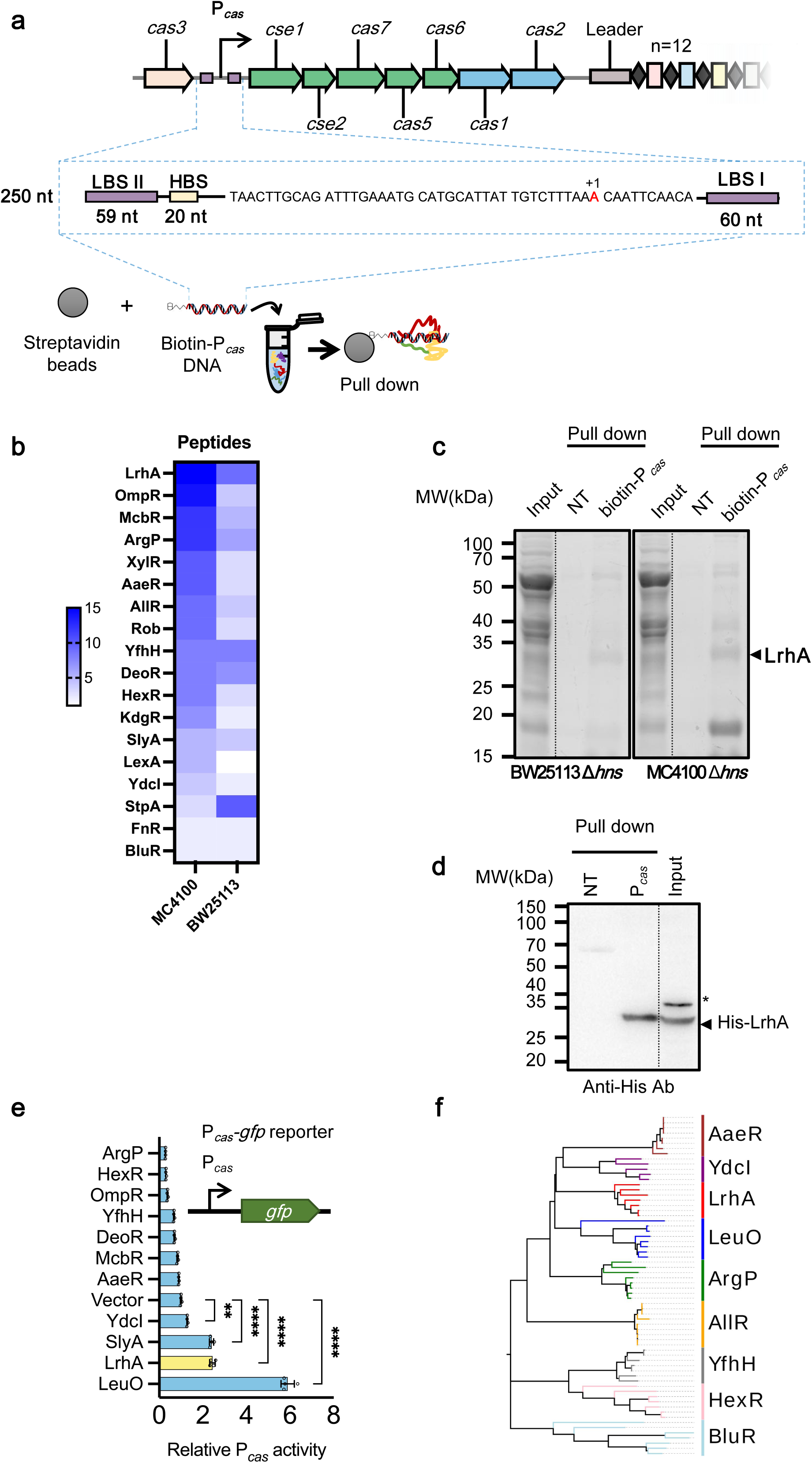
Screen for new regulators modulating transcription of *cas* genes. (**a**) Schematic illustration of the CRISPR-Cas locus in *E. coli*. Repeats (diamonds) in the CRISPR array are interspersed with 12 unique spacer sequences (rectangles). The promoter of the *cas* operon (P*_cas_*) is indicated by the black arrow. The P*_cas_* DNA probe for pull-down assay contains a transcription start site (red letter), LeuO binding site I and II (LBS I/II; purple bars), and H-NS binding site (HBS; yellow bar). For the pull-down proteomic experiment, a biotinylated probe (biotin-P*_cas_* DNA) was immobilized onto streptavidin beads, followed by incubation with extracts from *E. coli* MC4100 Δ*hns* (ZJUTCBB0015) or BW25113 Δ*hns* (ZJUTCBB0009) before SDS-PAGE electrophoresis. (**b**) A heatmap shows the number of peptides pulled down by P*_cas_* DNA probe. (**c**) SDS-PAGE gel electrophoresis of nontargeted biocytin (negative control, NT) or biotin-P*_cas_* DNA-binding proteins. NT or biotin-P*_cas_* samples were digested in gel with trypsin, and corresponding peptides were analyzed by mass spectrometry. An arrow denotes the band corresponding to LrhA (34.593 kDa). (**d**) Pull-down assay of physical interaction between LrhA and P*_cas_*. The biotinylated probe (biotin-P*_cas_* DNA) was immobilized onto streptavidin beads and incubated with His-LrhA from cell extracts before (lane 3, input) and after (lane 2, P*_cas_*) pull-down, followed by western blot assay with antibody against His-tag. Nontargeted biocytin (NT) was used as a negative control (lane 1). Arrow indicates the band of His-LrhA, and the asterisk indicates a non-specific band. (**e**) Effects of 12 predicted TFs and LeuO on the P*_cas_* activity in *E. coli* BW25113 Δ*hns* Δ*leuO* (EC73). The expression of TFs was induced by adding 10 mM arabinose. The P*_cas_* activity was evaluated by GFP/OD_600_ at 37 ℃, with the relative expression level of Δ*hns* Δ*leuO* carrying empty vector pSU19 (Vector) as 1 arbitrary unit. All bars represent the mean, and error bars denote standard deviation. Individual biological replicates are shown (*n* = 4). Statistical significance was determined using one-way ANOVA with Dunnett’s multiple comparisons test. ****, *P* < 0.0001; **, *P* < 0.01 (**f**) Phylogenetic analysis of 9 HTH-type P*_cas_*-binding TFs from 5-8 different bacteria species.

Remarkably, the LTTR LrhA was highly abundant in protein mixtures bound to P*_cas_* in both strains (Fig. 1b and c), with high coverage of peptide sequences (Fig. S3). Subsequent pull-down assay confirmed that purified LrhA with a 6×His-tag bound P*_cas_,* indicating that LrhA could be a potential regulator of the CRISPR-Cas system in *E. coli* (Fig. 1d).

To check whether these P*_cas_*-binding TFs regulate transcription of the *cas* operon, their effects on the activity of P*_cas_* were evaluated with a previously described fluorescence reporter system^64^ (Fig. 1e). To avoid the effect of the known strong regulators H-NS and LeuO, the 12 predicted TFs, and LeuO (as the positive control), were separately over-expressed in a Δ*hns* Δ*leuO* mutant. SlyA promoted the activity of P*_cas_* by 2.4-fold (Fig. 1e), in agreement with the previous report ^67^. Eight TFs contain the HTH-type DNA binding domain, which is also present in LeuO (Fig. 1f). The LTTR ArgP, known to act either as a transcriptional activator or a repressor depending on its binding with two effectors lysine and arginine^68^, reduced the activity of P*_cas_* by 70%, while LrhA increased the activity of P*_cas_* by more than 2-fold (Fig. 1e). Although YdcI and AaeR also belong to LTTRs and bind to P*_cas_*, they showed minimal (if any) effect on the activity of P*_cas_* (Fig. 1e). Thus, we narrowed down to the regulators LrhA, SlyA, ArgP and HexR below. To investigate their potential functions on regulating CRISPR interference, a conjugative plasmid pTc, that carries a DNA fragment with four protospacer-adjacent motif (PAM)-containing spacers from pT^64^, was generated. By examining clearance of the CRISPR-targeted plasmid (pTc) and the CRISPR-non-targeted plasmid (pNTc) in the BW25113 Δ*hns* Δ*leuO* mutant during conjugation. As shown in Fig. S4, LrhA and LeuO decreased the conjugation frequencies of pTc but not pNTc after 24h growth, exhibiting a higher CRISPR immunity than the Vector. ArgP did not affect the clearance rates of both plasmids significantly, with its CRISPR immunity unaffected. Overexpression of HexR or SlyA reduced plasmid stability, thus the conjugation frequencies of both plasmids were not detected (Fig. S4). Therefore, we focused on LrhA in the following investigation.

### LrhA activates transcription of the *cas* operon by binding to P*_cas_*

We investigated transcriptional regulation of P*_cas_* in *E. coli* K12 BW25113, which has an active type I-E CRIPSR-Cas system^65^, and ER2738, an F^+^ *E. coli* strain which is commonly used as a host strain for M13 phage amplification. ER2738 possesses not only intact *hns* and *leuO* genes (GenBank accession number PP194455, PP194456, PP112993), but also *cas* genes and CRISPR arrays, and their promoters (Fig. S5, GenBank accession number, PP194457), sequences of which are identical to that in BW25113. By using the P*_cas_*-*gfp* reporter, we observed that the P*_cas_* activity was 2.5-fold lower in ER2738 WT than in ER2738 Δ*hns*, while P*_cas_* activity was 14.5-fold lower in BW25113 WT than in BW25113 Δ*hns* (Fig. 2a), indicating that the repression effect of H-NS on P*_cas_* is remarkably weaker in ER2738 than in BW25113. qPCR analysis revealed that the amount of *hns* mRNA was more than 10-fold lower in ER2738 than in BW25113, indicating low expression of H-NS in the former strain (Fig. 2b). We inactivated *lrhA* in the Δ*hns* and Δ*hns* Δ*leuO* mutants to further evaluated how LrhA controls the transcription of the *cas* operon in both ER2738 and BW25113. In agreement with the previous report that LrhA represses transcription of the motility master regulator operon *flhDC*^69^, deleting *lrhA* increased the diameter of swarm rings (Fig. S6), showing correct construction of these strains. We detected no significant effect of chromosomally expressed LrhA on the transcription of the first gene of the *cas* operon (i.e., *cse1*) in BW25113 WT (Fig. 2c), while transcription of *cse1* was 4.4-fold lower in ER2738 Δ*lrhA* than in ER2738 WT (Fig. 2d), indicating that LrhA displays stronger effect on P*_cas_* in ER2738 than in BW25113. In the BW25113 Δ*hns* mutant, deleting *lrhA* decreased transcription of *cse1* by 11%. Further deleting *leuO* made LrhA exhibit a more pronounced effect on the activity of P*_cas_*. Deleting *lrhA* decreased transcription of *cse1* by 26% in the BW25113 Δ*hns* Δ*leuO* mutant (Fig. 2c). Complementing *lrhA* restored transcription of *cse1* in BW25113 Δ*hns* Δ*leuO* mutants (Fig. S7), confirming that LrhA is able to activate P*_cas_* (Fig. 2e). In contrast to moderate activation of the *cas* operon by LrhA, LeuO exhibited a stronger stimulation effect on P*_cas_*. In both BW25113 Δ*hns* and Δ*hns* Δ*lrhA* mutants, deleting *leuO* decreased transcription of *cse1* by more than 60% (Fig. 2c).

**Figure 2.**
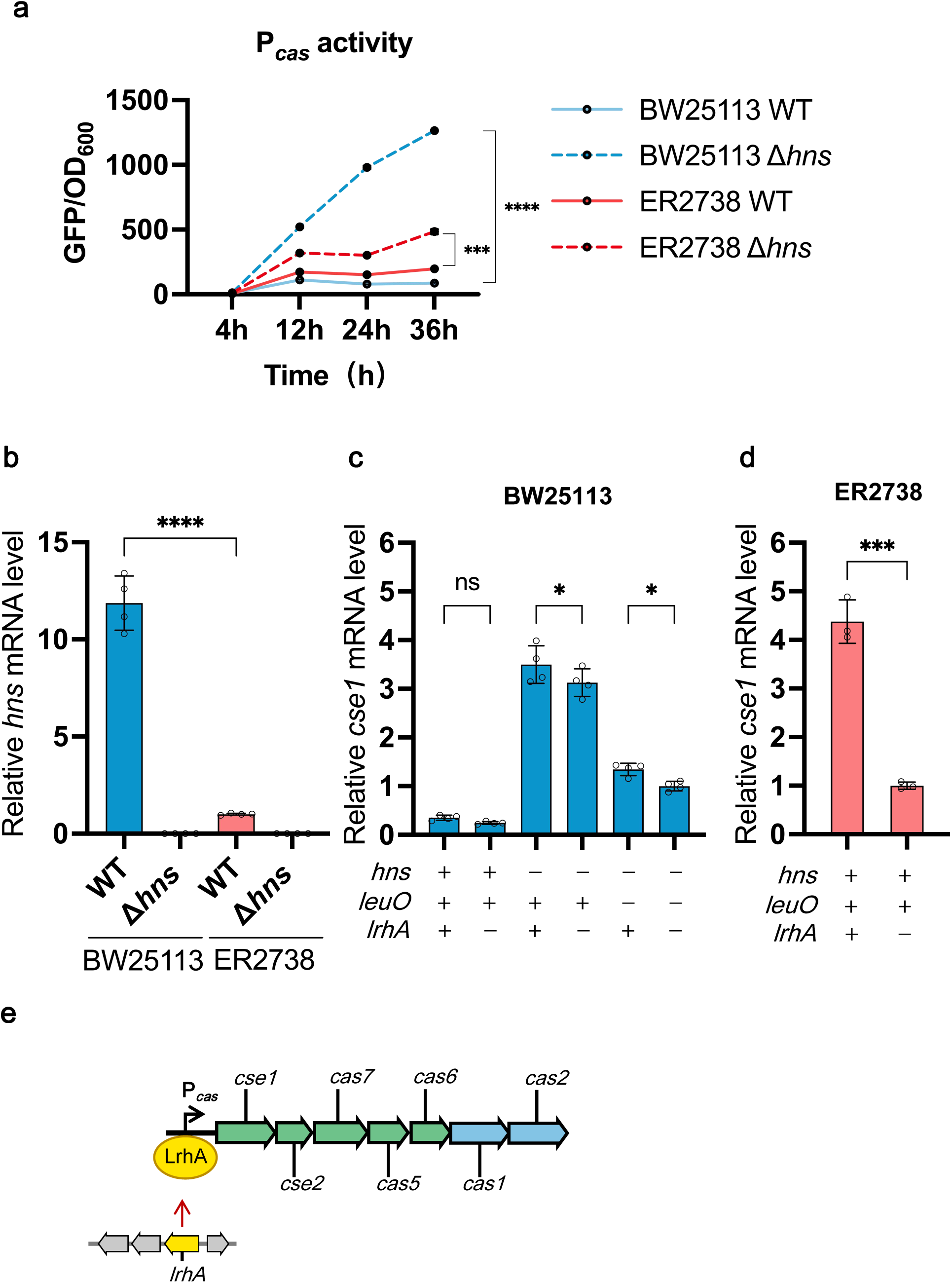
LrhA activates transcription of the *cas* operon by binding to P*_cas_*. (**a**) The P*_cas_* activity of BW25113 WT (blue solid line), BW25113 Δ*hns* (EC71) (blue dashed line), ER2738 WT (red solid line) and ER2738 Δ*hns* (EC74) (red dashed line) cells grown at 37 ℃, which was evaluated by GFP/OD_600_. Statistical significance was assessed using a two-way ANOVA (*n* = 3). ****, *P* < 0.0001; ***, *P* < 0.001. (**b**) qPCR analysis of *hns* in BW25113 WT, BW25113 Δ*hns* (EC71), ER2738 WT and ER2738 Δ*hns* (EC94) cells grown at 37 ℃ for 36 h, with the relative expression level of ER2738 Δ*hns* as 1 arbitrary unit. Statistical significance was assessed using a one-way ANOVA. Bars are the means and error bars ± SD. ****, *P* < 0.0001. Individual biological replicates are shown (*n* = 4). (**c**) Effects of genomic *lrhA* on the P*_cas_* activity (indicated by *cse1*) assessed by qPCR assay. P*_cas_* activities were compared in BW25113 WT, Δ*lrhA* (EC75), Δ*hns* (EC71), Δ*hns* Δ*lrhA* (EC76) strain, Δ*hns* Δ*leuO* (EC73), and BW25113 Δ*hns* Δ*leuO* Δ*lrhA* (EC77) grown in LB medium at 37 ℃ for 36 h, with the relative expression level of BW25113 Δ*hns* Δ*leuO* Δ*lrhA* as 1 arbitrary unit. Statistical significance was assessed using a one-way ANOVA. *, *P* < 0.05. Individual biological replicates are shown (*n* = 4). (**d**) qPCR analysis of *cse1* in ER2738 WT and ER2738 Δ*lrhA* (EC87) cells grown at 37 ℃, with the relative expression level of ER2738 Δ*lrhA* as 1 arbitrary unit. Statistical significance was assessed using a two-tailed Student’ s t-test. ***, *P* < 0.001. Individual biological replicates are shown (*n* = 3). (**e**) Schematic depiction of the regulation of P*_cas_* by LrhA.

The LrhA-binding site was reported to partially overlap with a LeuO-binding site on the promoter of *leuO*^70^. Previous work has identified two LeuO binding sites (LBS I and LBS II) within P*_cas_* ^60, 61^ (Fig. S1). The observation that the presence of LeuO interfered with the regulation of P*_cas_* by LrhA prompted us to consider whether LrhA and LeuO compete for the same binding sites on P*_cas_*. To check this, DNA probes lacking LBS I and/or LBS II were generated (named ‘P*_cas_* del LBS I’, ‘P*_cas_* del LBS II’, and ‘P*_cas_* short probe’) and EMSA was employed to determine the effect of LBS I and LBS II on the efficiency of LrhA binding to P*_cas_*. As shown in Fig. 3a, 700 nM of His-LrhA was sufficient to retard the mobility of the full-length P*_cas_* probe (P*_cas_* FL). However, deleting LBS I resulted in the abolishment of LrhA binding with P*_cas_*, even the concentration of LrhA reached 1400 nM, while deleting LBS II only partially impaired LrhA binding with P*_cas_*. (Fig. 3a). These observations support that LrhA binds to both LBS I and LBS II.

**Figure 3.**
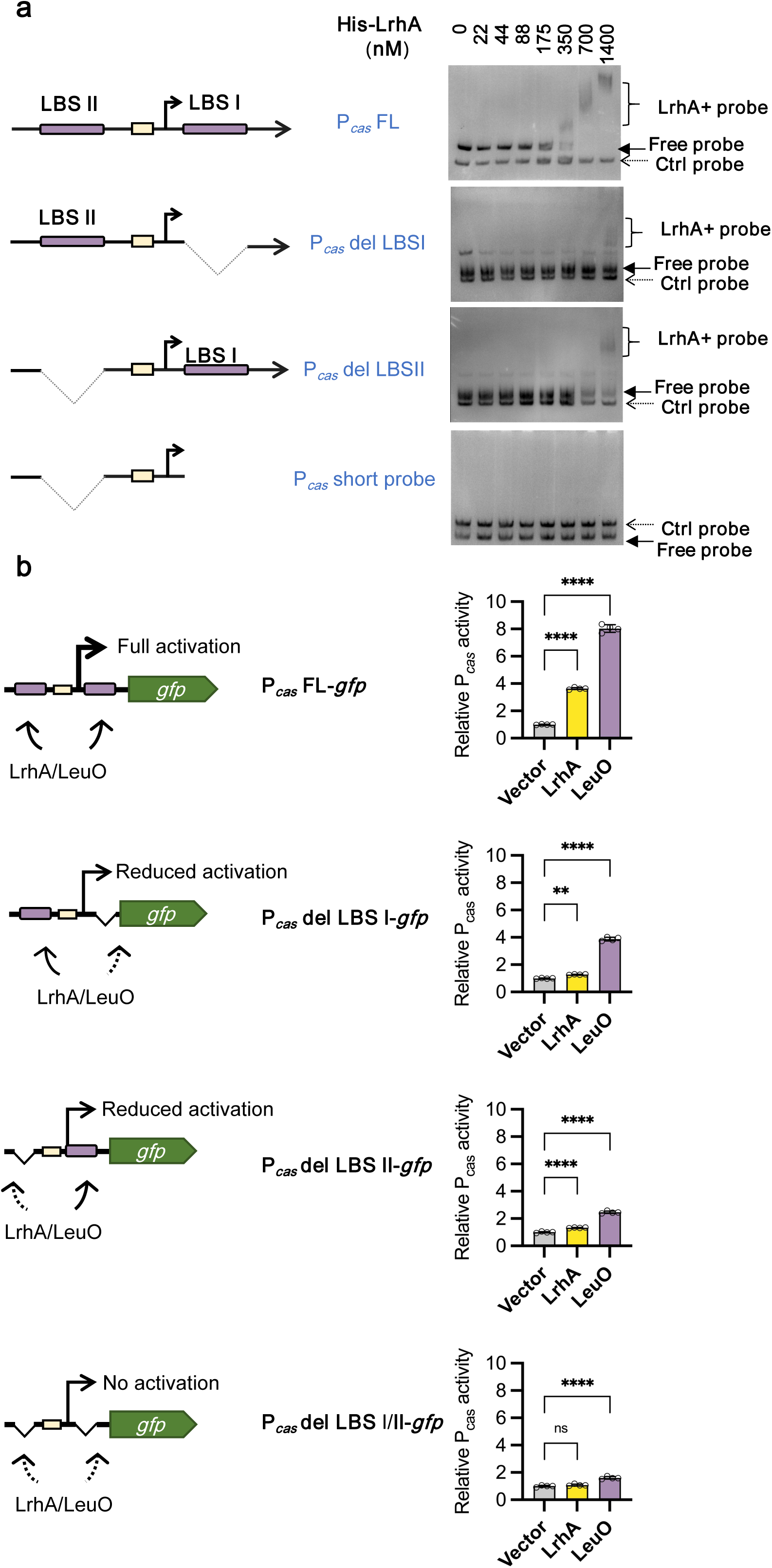
Identification of LrhA binding sites on the promoter of the *cas* operon. (**a**) (**left**) Schematic illustration of the P*_cas_* region for truncation analysis. LBS I and LBS II indicate two DNA-binding sites of LeuO located upstream and downstream of the transcriptional start site (TSS, black arrow) on P*_cas_* (purple bars). HBS indicates the DNA-binding site of the H-NS (yellow bar). (**right**) electrophoretic mobility shift (EMSA) assays of full-length or truncated P*_cas_* DNA fragments by LrhA. The P*_cas_* probes including full-length P*_cas_* (P*_cas_* FL), P*_cas_* with LBS I deleted (P*_cas_* del LBS I), P*_cas_* with LBS II deleted (P*_cas_* del LBS II) and P*_cas_* with both LBS I and LBS II deleted (P*_cas_* short probe) were generated. 1 nM P*_cas_* probes were separately incubated with a serial concentration of purified His-LrhA before EMSA assays with 5% native polyacrylamide gels (solid arrow for free P*_cas_* probes, and braces for LrhA+probe complex). A 175 bp probe of *lacZα* fragment was used as the control (Ctrl probe, dashed arrow). (**b**) (**left**) Schematic illustration of DNA binding sites required for activating P*_cas_* by LrhA and LeuO. Transcriptional activities of full-length P*_cas_*, P*_cas_* del LBS I, P*_cas_* del LBS II and P*_cas_* del LBS I/II were compared by using the P*_cas-_gfp* reporter. (**right**) Effects of disruption of LBS I and LBS II on activation of P*_cas_* by LrhA. The P*_cas_* activity was evaluated by GFP/OD_600_, with the relative expression level of BW25113 Δ*hns* Δ*leuO* carrying the empty vector as 1 arbitrary unit. Bars are the means and error bars ± SD. Individual biological replicates are shown (*n* = 4). Statistical significance was determined using one-way ANOVA with Dunnett’s multiple comparisons test. ****, *P* < 0.0001; **, *P* < 0.01.

To know whether LBS I and LBS II were essential to activate P*_cas_* by LrhA and LeuO *in vivo*, we generated P*_cas_* containing disrupted LBS I and/or LBS II (Fig. 3b), and compared *in vivo* activities of disrupted P*_cas_* with that of P*_cas_* FL in response to expression of LrhA and LeuO. Over-expressing LrhA and LeuO increased the transcriptional activity of P*_cas_* FL by more than 3- and 8-fold, respectively (Fig. 3b). In contrast, the activity of P*_cas_* del LBS I in the BW25113 Δ*hns* Δ*leuO* mutant expressing LrhA or LeuO was decreased to the background level or 3.88-fold higher than the background level, respectively (Fig. 3b). This observation indicates that LBS I is indispensable for activation of P*_cas_* by LrhA, and essential for full stimulation of P*_cas_* by LeuO. Interestingly, the activity of P*_cas_* del LBS II was also remarkably reduced in the BW25113 Δ*hns* Δ*leuO* mutant expressing either LrhA or LeuO (Fig. 3b), revealing that LBS II is required for activation of P*_cas_* by both LrhA and LeuO. As expected, when both LBS I and LBS II were deleted, the expression of LrhA or LeuO no longer stimulated the expression of the impaired P*_cas_* (Fig. 3b). Taken together, these data clearly show that both LBS I and LBS II are essential for activation of P*_cas_* by LrhA and LeuO *in vivo*.

### LrhA activates CRISPR interference against M13 phage infection

Having established LrhA as a regulator of P*_cas_*, we sought to test whether it could play a role in regulating CRISPR interference against bacteriophage in the wild-type *E. coli* ER2738, which contains an F-factor encoding pili that are required for M13 phage adsorption ^71^. A spacer (g8 spacer) matching the gene 8 of M13 phage was integrated into the CRISPR I array of the wild-type ER2738 strain (WT-g8), which was expected to confer resistance to M13 phage infection (Fig. 4a). The ER2738 Δ*lrhA* mutant showed swarm rings with increased diameter compared with ER2738 WT (Fig. S6c). Inactivation of *lrhA* in ER2738-g8 (Δ*lrhA*-g8) reduced the efficiency of plaquing (EOP) by ∼10^6^-fold, while complementing *lrhA* in the *ΔlrhA* mutant (*lrhA*^c^-g8) fully restored its resistance to M13 phage (Fig. 4b), revealing that LrhA expressed from the genome strongly activates the CRISPR-Cas immunity against infection of the phage. Inactivation of *cas3* abolished the CRISPR immunity in WT-g8 (Fig. S8). Together, these results demonstrate that LrhA plays a crucial role in promoting the CRISPR interference against phage infection in a wild-type *E. coli* strain (Fig. 4c).

**Figure 4.**
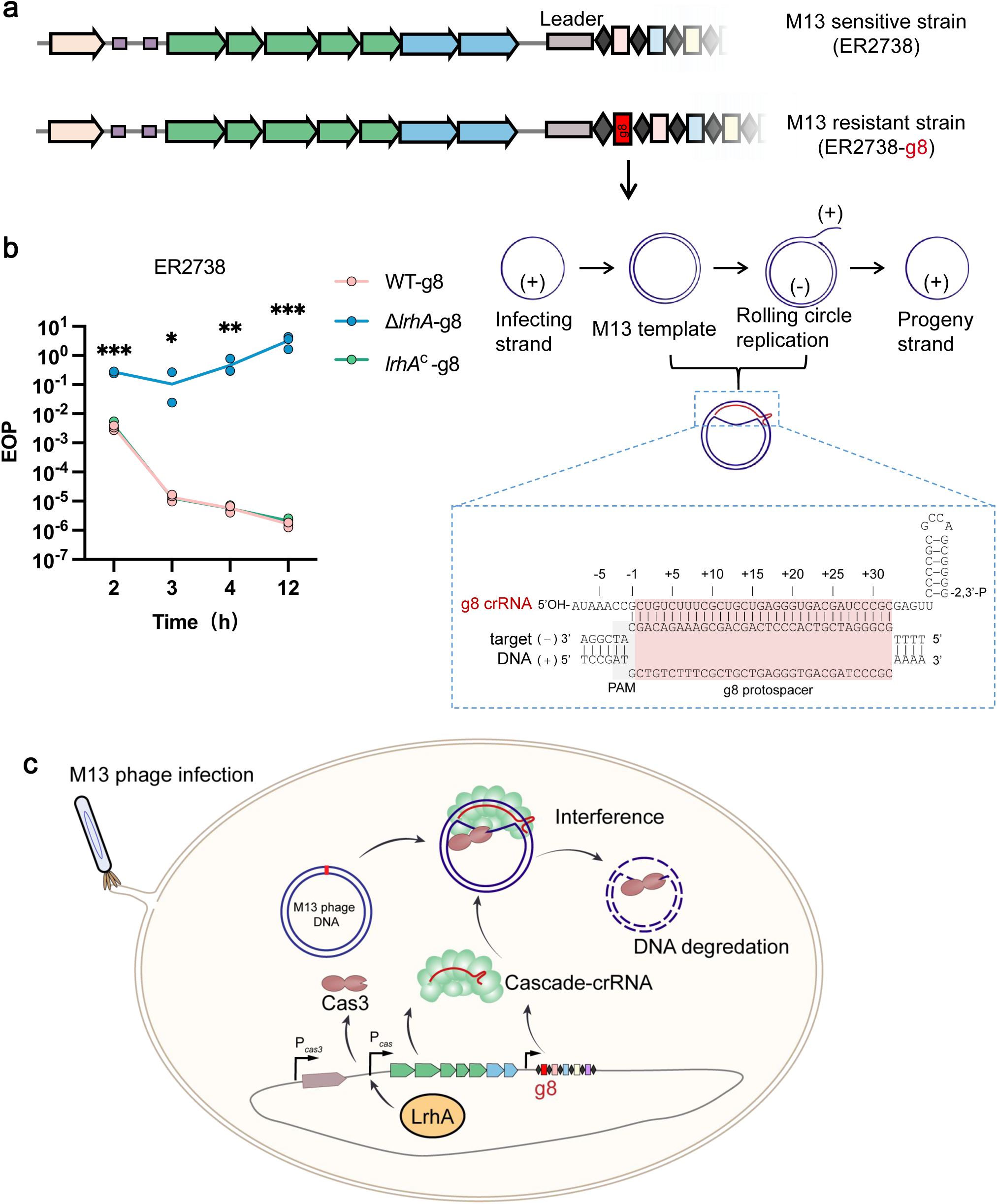
LrhA activates CRISPR-Cas immunity against M13 phage. (**a**) Schematic illustration of the engineered CRISPR cassette carrying a g8 spacer in the genomic CRISPR I locus. In M13 targeting cells, CRISPR I contains an additional g8 spacer (red rectangle), conferring cells with resistance to M13 phage infection. During infection, phage DNA enters the cell as a circular (+) single-stranded (ss) DNA, followed by the formation of double-stranded (ds) phage genome DNA (template), which is replicated in a rolling circle form to generate progeny (+) strand that is packaged in mature phage. M13 phage DNA in the dsDNA form (i.e., template and rolling circle replication) can be recognized by g8 crRNA-guided Cascade complex. (**b**) Quantification of the efficiency of plating (EOP) at 2-12 h after infection with M13 phage. Efficiency of plaquing (EOP) was calculated as a ratio of phage titers observed on cells with g8 spacer (WT-g8, EC89; Δ*lrhA*-g8, EC90 and *lrhAc*-g8, EC91) to that on cells without g8 spacer (non-targeting cells, WT, ER2738; Δ*lrhA*, EC87 and *lrhA^c^*, EC88). Individual biological replicates are shown (*n* = 3). Statistical significance between WT-g8 and Δ*lrhA*-g8 at 2-12h was determined using multiple t-tests. *, *P* < 0.05; **, *P* < 0.01; ***, *P* < 0.001 (**c**) Schematic illustration of regulation of the CRISPR-Cas system against M13 phage infection by LrhA.

### LrhA promotes clearance of the CRISPR-targeted plasmid

We examined the role of LrhA on transferring and transferred CRISPR-targeted plasmids. Although deletion of *lrhA* showed no effect on CRISPR immunity against pTc during conjugation (Fig. S9a-c), we observed that LrhA promoted CRISPR immunity against the transferred CRISPR-targeted plasmid in *E. coli* BW25113 lacking *hns* and/or *leuO*. The CRISPR immunity against transferred plasmid was assessed by evaluating the clearance rate of CRISPR-targeted plasmid in *lrhA^+^*and *lrhA*^−^ strains. The relative CRISPR immunity in the *lrhA^+^*strain was ∼3.5-fold of that in the *lrhA*^−^strain with deletion of *hns* (Fig. 5a-b) and ∼9.8-fold of that in the *lrhA*^−^ strain with further deletion of *leuO* (Fig. 5b), demonstrating that LrhA activates type I-E CRISPR-Cas system for clearing the CRISPR-targeted plasmid independent upon H-NS and LeuO. Given that deleting *leuO* led LrhA to exhibit a stronger effect on promoting clearance of CRISPR-targeted plasmid, promotion of the CRISPR immunity by LrhA could be partially masked by LeuO.

**Figure 5.**
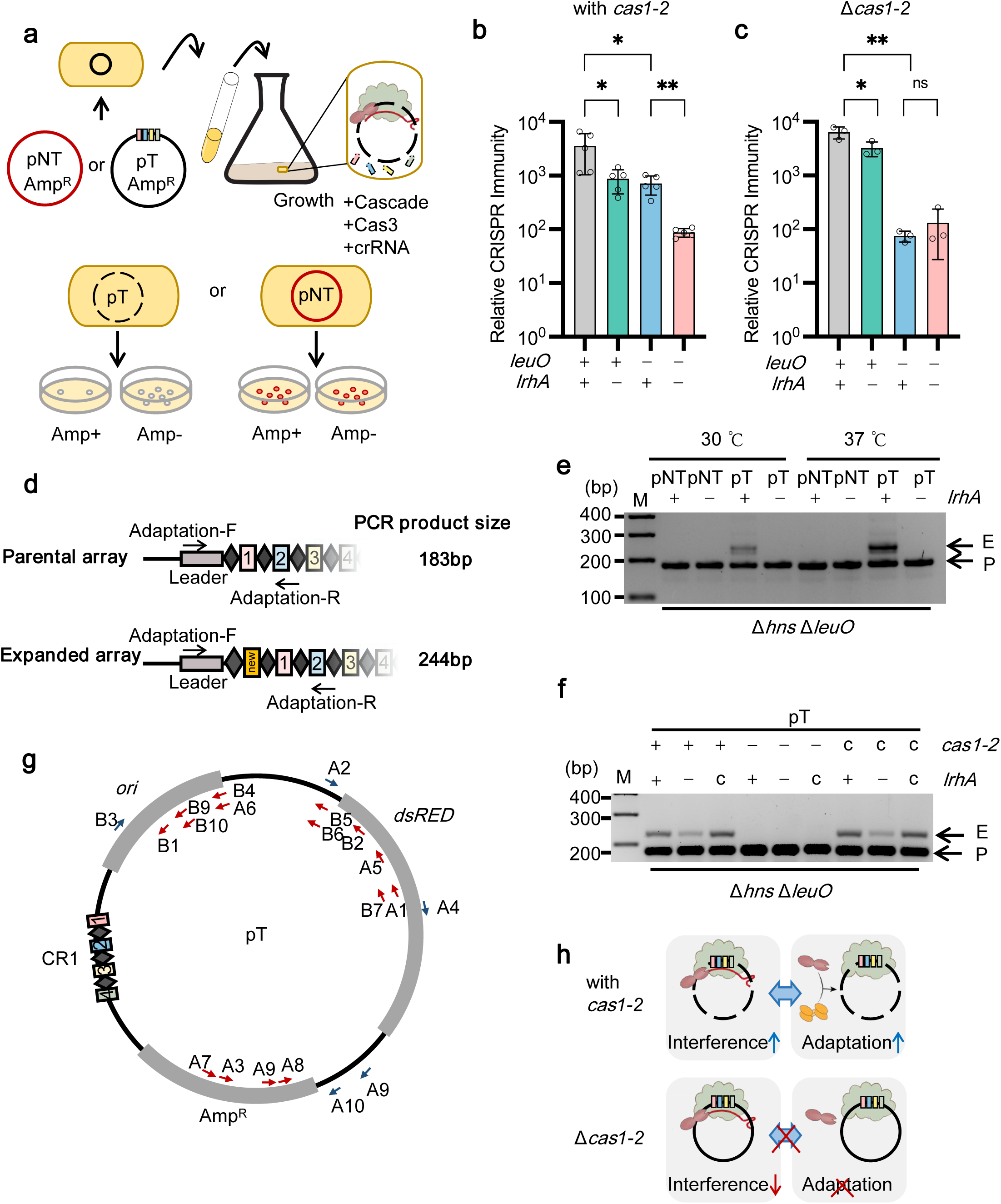
LrhA activates both CRISPR-Cas interference and adaptation. (**a**) Schematic illustration for evaluating the CRISPR-Cas interference activity by plasmid loss assay. (**b** and **c**) Relative CRISPR immunity assessed by plasmid loss assay in *E. coli* strains BW25113 Δ*hns* (EC71), Δ*hns* Δ*lrhA* (EC76), Δ*hns* Δ*leuO* (*lrhA*^+^, EC73) and Δ*hns* Δ*leuO* Δ*lrhA* (*lrhA*^−^, EC77) grown in antibiotic-free LB medium for 36 h. Statistical significance was assessed using a two-tailed Student’s t-test. Bars are the means and error bars ± SD. Individual biological replicates are shown (*n* = 5 for *b* and *n* = 3 for c). *, *P* < 0.05; **, *P* < 0.01. (**d**) Schematics of the spacer acquisition assay in (**e** and **f**). During the adaptation process, a new spacer (a yellow rectangle) can be inserted into the leader end of the CRISPR array. PCR was performed to amplify the leader-spacer region using the primers matching the leader and the second spacer, which were indicated as flagged arrows. (**e**) Spacer-acquisition assay by PCR amplification of BW25113 Δ*hns*Δ*leuO* (*lrhA*^+^, EC73), BW25113 Δ*hns* Δ*leuO* Δ*lrhA* (*lrhA*^−^, EC77) cells carrying pT or pNT plasmids, which were grown at 37 ℃ or 30 ℃ for 36 h. PCR products were electrophoresed on a 2% agarose gel and imaged. Parental (P) and expanded bands (E) are indicated on the right while DNA marker (M) positions are displayed on the left. Gels are representative of two experiments yielding similar results. (**f**) spacer-acquisition assay of *lrhA*^+^, *lrhA*^−^ and *lrhA*^c^ with *cas1*-*cas2* deletion (Δ*cas1-2,* EC81, EC82, EC83) or its complementation (*cas1-2* ^c^, EC84, EC85, EC86). (**g**) Mapping of newly formed spacers on pT plasmid DNA. The DNA of the expanded array was sequenced by ligating purified PCR products into T vectors. The position and orientation of the sequenced spacers are marked on the pT plasmid map (blue arrows pointing clockwise and red arrows pointing counterclockwise). Protospacers are numbered according to the strain from which the spacers were sequenced, as listed in Table S3. CR1 indicates 4 spacers targeted by the type I-E CRISPR-Cas system of *E. coli*. (**h**) Schematic illustration of the relationship between CRISPR interference and adaptation regulated by LrhA. In the presence of *cas1-2*, LrhA promotes the positive feedback circuit between CRISPR interference and adaptation (upper panel). In the absence of *cas1-2*, LrhA shows a weak effect on interference (lower panel).

### Promoting primed adaptation by LrhA accelerates clearance of the CRISPR-targeted plasmid

We observed an apparent effect of LrhA on CRISPR immunity against transferred plasmid during long-term growth but not transferring plasmid during conjugation. Given that weak interference often allows DNA to persist longer and promotes acquisition of new spacers that allows bacteria to mount more efficient CRISPR interference ^32, 34, 72^, we hypothesized that intermediate activation of the *cas* operon by LrhA might promote clearance of pT through better coordinating interference and primed adaptation. To test this hypothesis, we monitored CRISPR array expansion in response to pT and pNT in *lrhA^+^*and *lrhA*^−^ strains (Fig. 5d). Compared with that in the *lrhA^+^* strain, fewer expansion events were observed in the *lrhA*^−^ mutant carrying pT after 36 h of incubation at 37 ℃, showing that LrhA is required for efficient adaptation (Fig.5e, Fig. S10). Complementing *lrhA* on the genome of the *lrhA*^−^ mutant (*lrhA*^c^) restored adaptation (Fig. 5f). In contrast, CRISPR array expansion was not observed in the *lrhA^+^* and *lrhA*^−^strains carrying pNT (Fig. 5e). Sequencing analysis of the expansion bands from *lrhA^+^*strains carrying pT revealed that all newly formed spacers (20/20) were derived from the CRISPR targeted plasmid (Fig. 5g), with a strong preference for an AAG PAM (16/20) during adaptation, and a pronounced bias in the DNA strand (15/20) matching the orientation of the CRISPR-targeting units (CR1) (Table S3), in agreement with typical traits of primed CRISPR adaptation^32, 34^. Compared with other regions on the CRISPR-targeted plasmid, more protospacers were mapped to encoding genes (i.e., *bla* and *dsRED*) and the replication origin (Fig. 5g).

To confirm that LrhA promoted clearance of pT through reinforcing the interplay between CRISPR-Cas interference and primed adaptation, we deleted both *cas1* and *cas2* (Δ*cas1-2*) in *lrhA^+^*, *lrhA*^−^ and *lrhA*^c^ strains, and examined the CRISPR interference in these strains through the plasmid loss assay. Deletion of *cas1*-*cas2* abolished the CRISPR array expansion (Fig. 5f), and reduced the effect of LrhA on CRISPR immunity against plasmid (Fig. 5c, Fig S11), revealing that LrhA activates CRISPR immunity through promoting the positive feedback of interference and adaptation processes (Fig. 5h). Loss of CRISPR expansion from Δ*cas1-2* cells was restored by genomic complementation of *cas1*-*cas2* (*cas1-2*^c^) (Fig. 5f), consistent with critical role of *cas1*-*cas2* in adaption. Activation of CRISPR interference by LrhA was regained by complementation of *cas1*-*cas2* (Fig. S11), confirming that Cas1-Cas2 was essential to LrhA-regulated adaptation-mediated interference.

## DISCUSSION

In this study, we have addressed the important question of how the CRISPR-Cas system is regulated through coordinating interference and adaptation. By combining DNA pull-down and mass spectrometry, we identified a novel regulator, LrhA, that enhances immunity against phage infections by directly blocking the entry of phage DNA and facilitates a reciprocal interplay between interference and adaptation, thereby providing protection against horizontally transferred plasmids in *E. coli*. Our study not only provides new insights into how bacteria control the adaptive immunity against exogenous DNA under different scenarios, adding a new layer of regulatory complexity in bacterial defense, but also introduces a functional approach to screen any transcriptional regulators that directly bind to promoters in bacteria.

Early reports suggested that primed adaptation was triggered by suboptimal Cascade-protospacer interactions that could lead to weak interference^32, 58, 73^, whereas others indicated that optimal Cascade-protospacer interactions also led to both interference and primed adaptation, which was fueled by interference^35, 53, 72, 74^. Evidence remains lacking for intracellular regulation of CRISPR immunity through coordination of interference and adaptation. In this study, we found that LrhA moderately stimulated transcription of *cas* genes but strongly promoted the clearance of CRISPR-targeted plasmid. Adaptation occurred more frequently in the *lrhA*^+^ *E. coli* strains harboring the CRISPR-targeted plasmid (Fig. 5, and Fig. S10), revealing that LrhA can promote interference-driven spacer acquisition. Given that inactivation of *cas1* and *cas2* remarkably reduced LrhA-stimulated CRISPR immunity (Fig. 5c), we propose that moderate activation of transcription of *cas* genes by LrhA enables the positive feedback circuit between CRISPR interference and adaptation: cleavage of the CRISPR-targeted plasmid produces more short DNA fragments for accelerating primed adaptation, which in turn enhances clearance of the CRISPR-targeted plasmid (Fig. S12). In contrast, strong activation of transcription of *cas* genes by LeuO was unlikely to trigger the positive feedback circuit, and exhibited weaker effect on clearance of the CRISPR-targeted plasmid. Analysis of the acquired spacers revealed an unequal distribution (Fig. 5g), suggesting that certain sequences could be preferentially integrated as new spacers.

The transcriptional regulators of type I-E CRISPR-Cas system have not been screened thoroughly. In a previous study of identification of the potential host factors involved in adaptation, Yoganand *et al* employed the CRISPR/dCas9 mediated immunoprecipitation (IP) to identify host factors that bound the CRISPR leader region^75^. Compared to the reported CRISPR/dCas9-mediated IP screening strategy, the pull-down assay employed in this study is significantly simpler and facilitates the identification of a larger number of potential proteins, owing to the ease of obtaining sufficient amounts of biotin-labeled probes, albeit at the expense of increased false positive detection. Nevertheless, subsequent screening with a fluorescence reporter helps to the presence of unwanted proteins. By using the DNA pull-down assay with a DNA probe containing P*_cas_* as the bait, we screened out a number of host factors that associated with P*_cas_* from the cell lysate. Particularly, both our *in vivo* and *in vitro* data reveal that the pleiotropic LTTRs LrhA activate the *cas* operon by binding to the LeuO binding sites in P*_cas_*. LTTR family proteins seem to extensively regulate diverse types of CRISPR-Cas systems in different bacteria. LrhA (named PigU in *Serratia*) was reported to suppress the type III and type I-F but not the type I-E CRISRP-Cas system^76^, indicating that this regulator plays differential roles in different bacteria. In *Salmonella enterica*, transcription of the CRISPR-Cas system was by two LTTRs with opposing functions (i.e., LeuO and LRP) ^77^. LeuO is normally considered as an antagonist of H-NS, and activates gene transcription by relieving transcriptional suppression by H-NS, we found that both LrhA and LeuO promoted the activity of P*_cas_* in the Δ*hns* Δ*leuO* mutants (Fig. 1e, Fig. 2c), indicating that they both regulate the *cas* operon independent of H-NS.

Although both LrhA and LeuO bind P*_cas_*, their impacts on regulation of the type I-E CRISPR-Cas system and other physiological functions are context-dependent. LrhA strongly stimulated the CRISPR-Cas activity against M13 phage in the wild-type *E. coli* ER2738 (Fig. 4b), demonstrating that LrhA is capable of playing a dominant role in regulating CRISPR-Cas activity in *E. coli* upon phage infection. Roles of LeuO and LrhA are also different in regulating the CRISPR-Cas activity for clearing horizontally transferred plasmid. Although LeuO stimulated stronger transcription of *cas* genes, LrhA stimulated higher adaptive immunity in clearing CRISPR-targeted plasmid through better-coordinating interference and adaptation. The two regulators show different binding site dependencies at P*_cas_*. Apart from regulating the CRISPR-Cas system, LrhA and LeuO showed remarkable differences in controlling other physiological functions, such as the flagellar *flhDC* operon, which can be inhibited by LrhA, but not LeuO (Fig. S6). Moreover, LeuO activated transcription of the *cas* operon through direct interacting with the α subunit of the RNA polymerase (RpoA) on P*_cas_*, but the regulation of P*_cas_* by LrhA seemed independent of RpoA (unpublished data). Phylogenetic analysis revealed divergent evolution of LrhA and LeuO, which could belong to distinct subfamilies and separately evolve in bacteria (Fig. S13 and S14).The expression of LeuO can be activated by the heterodimer BglJ-RcsB^78^, Cyanase regulator CynR^70^, and antibiotic responses regulator YdcI^70^, while the expression of LrhA is stimulated by acid resistance-related gene *hdeD* ^79^. Hence, the regulation of the type I-E CRISPR-Cas system by LrhA and LeuO could have been developed in response to different environmental cues.

## MATERIALS AND METHODS

### Bacterial strains, plasmids, primers, growth conditions, and media

Bacterial strains, plasmids, and primers used in this study are listed in *Supplementary information* Table S4, Table S5 and Table S6. *E. coli* gene-deletion mutants were constructed by using a λ-RED recombination system expressed by the temperature-sensitive plasmid pKD46 ^80^. Corresponding mutants were examined through PCR. Recombinant plasmids expressing LrhA and other 10 TFs were constructed with a Transfer PCR method ^81^. Plasmids were isolated and purified with a plasmid isolation kit according to the manufacturer’s protocol (Axygen Biotech Co., Ltd.). *E. coli* cultures were grown at 37 ℃ in LB broth medium containing 1% (wt/vol) tryptone, 0.5% (wt/vol) yeast extract, and 1% (wt/vol) NaCl or on LB agar plates containing 1.5% (wt/vol) agar. When required, media were supplemented with ampicillin (100 μg mL^−1^), kanamycin (50 μg mL^−1^), or chloramphenicol (25 μg mL^−1^). Arabinose was added for inducing gene expression in liquid cultures where indicated. Cell growth was measured in a spectrophotometer (MD SpectraMax iD5 or Biotek Synergy 2). All experiments were repeated independently at least three times unless otherwise indicated.

### Pull-down assay

P*_cas_*-binding proteins were pulled-down with the streptavidin beads (Invitrogen Dynabeads™ M-270, 65305) according to the manufacturer’s protocol. (see “Pull-down assay” in *Supplementary information* Supplementary Methods).

### Mass spectrometry analysis

Mass spectrometry and analysis of the data were conducted by Micrometer Biotech Company. (see “Mass spectrometry analysis” in *Supplementary information* Supplementary Methods).

### Quantification of gene transcription

For quantifying P*_cas_* with reporter assay, plasmid pGLO-P*_cas_*-*gfp* was used for monitoring the transcription of the *cas* operon, by measuring the intensity of GFP fluorescence and cell optical density (OD_600_) on a microplate reader (MD, SpectraMax iD5), with a method that has previously been described^64^. When evaluating the GFP/OD_600_ level or the interference/adaptation effect, cells were grown in M9 minimal medium containing 61 mM K_2_HPO_4_·3H_2_O, 38.21 mM KH_2_PO_4_, 2.5 mM MgSO_4_, 15.14 mM (NH_4_)_2_SO_4_, 0.1% (wt/vol) tryptone, supplemented with 1% (wt/vol) glucose as the carbon source.

For the quantitative real-time PCR (qPCR) assay, bacteria were harvested from 0.4-1mL culture by centrifugation. Total RNA was extracted by using an RNA Extraction Kit (AG21017, Accurate Biotechnology Co., Ltd). Removal of the genomic DNA and reverse transcription were performed by using Evo M-MLV RT Mix Kit (with gDNA clean reaction mix) (AG11728) according to the manufacturer’s instructions. SYBR Green Premix Pro Taq HS qPCR Kit (AG11701) was used for qPCR, and the cycle thresholds were determined using a BioRad CFX96 Real-Time System. *rpoD* was used as the internal control. The primers for *cse1* and *rpoD* are listed in Table S6.

### Purification of LrhA Protein

The LrhA protein was purified by Ni-NTA Resin (ThermoFisher, 88222), according to the manufacturer’s protocol. (see “Purification of LrhA Protein” in *Supplementary information* Supplementary Methods).

### Electrophoretic mobility shift assay (EMSA)

EMSA was performed by using the LightShift Chemiluminescent EMSA kit (ThermoFisher, 20148) according to the manual guide. For more details, see “Electrophoretic mobility shift assay” in *Supplementary information* Supplementary Methods.

### Plasmid loss assay

The *E. coli* strains harboring the CRISPR-targeted plasmid pT (p*dsRED*-CR1) or the CRISPR-non-targeted plasmid pNT (p*dsRED*) were grown overnight in LB culture supplemented with ampicillin (100 μg mL^−1^). The overnight grown culture was washed and inoculated at a ratio of 2% into antibiotic-free LB medium, followed by incubation at 37 ℃ with shaking. At intervals, the culture was serially diluted and spread on the LB plate supplemented with or without ampicillin (100 μg mL^−1^). The number of cells maintained the plasmid was evaluated by comparing colonies on the LB plate supplemented with or without ampicillin. The relative CRISPR immunity was defined as the proportion of cells containing the pNT plasmid relative to that containing the pT plasmid.

### Spacer acquisition assay

Newly acquired spacers were detected by PCR using primers pair Adaptation-F and Adaptation-R. (see “Spacer acquisition assay” in *Supplementary information* Supplementary Methods).

### M13 phage sensitivity assay

*E. coli* ER2738 strains with or without chromosomal g8 spacers were grown in LB medium supplemented with M13 phage with a titer of 10^6^ plaque-forming units (PFU) per mL at 37 ℃. PFU of M13 phages from these bacterial cultures were counted at the indicated time points through a documented method ^32, 48^. Briefly, the supernatants were diluted tenfold, then mixed with 100 μL ER2738 (OD_600_ ∼ 0.5). The mixture was spread in 1 mL 0.75 % LB top agar and overlaid on LB plates containing 0.2 mM IPTG (isopropyl-β-D-thiogalactoside) and 40 μg mL^−1^ X-Gal (5-bromo-4-chloro-3-indolyl-b-Dgalactopyranoside) for visualization of M13 phage which carries the *lacZα* gene. After incubation at 37 ℃ overnight, the plaque-forming units (PFU) on plates were counted. Efficiency of plaquing (EOP) was calculated as a ratio of phage titers observed on cells with g8 spacer to that on cells without g8 spacer (non-targeting cells).

## Supporting information

supplementary informations

## ACKNOWLEDGMENTS

We thank Prof. Shishen Du at Wuhan University and Dr. Yanmei Zhang at Hangzhou Medical College for scientific advice on this manuscript. We also thank Prof. Zhiyong Sun at Zhejiang University of Technology, Prof. Pengjun Zhang at Hangzhou Normal University, Dr. Minjia Shen at Université Paris-Saclay, Ms. Xia Yao at Zhejiang University of Technology, and all members of the Sun laboratory for critically reading the manuscript and helpful discussions. We also would like to express our sincere gratitude to the editor and three anonymous reviewers for their valuable comments, which have greatly improved the quality of the manuscript.

## Author Contributions

M.Fang, F. W. and D.S. conceived the study; M.Fang, N.L., M.G., M.Fei, Y.L., M.Y., F. W. and D.S. designed research; M.Fang, N.L., M.G., M.Fei, Y.L., and M.Y., performed research; M.Fang, N.L., M.G., M.Fei, Y.L., M.Y., and D.S. analyzed data; and M.Fang., and D.S. wrote the paper.

