## supplementary informations for "LrhA promotes CRISPR-Cas immunity by enhancing primed adaptation in *Escherichia coli*"

**This PDF file includes:**

Supplementary Methods

Figures S1 to S15

Tables S1 to S6

SI References

### SUPPLEMENTARY METHOTS

***Pull-down assay***

To prepare samples for mass spectrometric analysis of proteins binding to $P_{cas}$, 8 pmol biotinylated $P_{cas}$ DNA fragment was coated with 100 µL streptavidin beads (Invitrogen Dynabeads™ M-270, 65305) in the binding buffer A (10 mM Tris-HCl pH 7.5, 1 mM EDTA, 2 M NaCl) for 1 h at the room temperature. Biocytin, a derivative of biotin, was used as the negative control. From a 5 mL culture of *E. coli hns* deletion-mutant grown overnight at 37 ℃, proteins were extracted by sonication in the binding buffer B (10 mM Tris-HCl pH 7.5, 1 mM EDTA, 50 mM NaCl, 0.4 mM DTT and 10% glycerol), supplemented with protease inhibitor cocktail (Roche Co., Ltd.). The crude lysates were centrifuged at 15000×*g* for 10 minutes. The supernatant was collected, pre-cleared by empty streptavidin beads, and then incubated with 50 µL $P_{cas}$ DNA coated beads overnight at 4 °C under constant rotation (20 rpm). Beads were washed 5 times with the binding buffer B, followed by washing 3 times with the binding buffer B supplemented with 1 mg salmon sperm DNA, and finally washing 5 times with the binding buffer B to remove non-specific proteins. DNA binding proteins were eluted in SDS sample loading buffer and loaded onto 10% SDS-PAGE, followed by the mass-spectrometric analysis of the gel bands as described below.

***Mass spectrometry analysis***

After 37 ℃ overnight in-gel digestion of proteins binding to $P_{cas}$ with trypsin, peptides were loaded onto a home-made reversed-phase analytical column (15-cm length, 75 µm i.d.), and analyzed with a gradient from 5% to 34% solvent B (0.1% formic acid in 80% acetonitrile) over 40 min, 34% to 38% solvent B for 5 min, 38% to 90% solvent B for 10 min, and then holding at 90% solvent B for 10 min, at a constant flow rate of 300 nL/min on an EASY-nLC 1000 UPLC system. Peptides were subjected to NSI source followed by tandem mass spectrometry (MS/MS) in Thermo Scientific Orbitrap Fusion coupled online to the UPLC. Survey full-scan MS spectra (m/z 350-1800) were acquired in the Orbitrap with resolution r of 70000 at m/z 400. The mass spectrometry proteomics data have been deposited to the ProteomeXchange Consortium (http://proteomecentral.proteomexchange.org) via the iProX partner repository with the dataset identifier PXD036879.

***Purification of LrhA Protein***

The *lrhA gene* was amplified and cloned into the pET28a vector to generate pET28a-His-*lrhA*, which encodes the LrhA protein fused to an N-terminal His-tag. The pET28a-His-*lrhA* plasmid was transformed into *E. coli* BL21 (DE3). The overnight culture was inoculated into 100 mL LB medium at a ratio of 1:100. When the culture was grown at 37 ℃ to an $OD_{600}$ of 0.4, expression of LrhA was induced with 0.5 mM IPTG (Beyotime, ST098) at 37 ℃ for 2 h. Cells were centrifuged at 8000 rpm for 5 min at 4 ℃. The cell pellet was resuspended in

PBS containing protease inhibitor and sonicated, and the lysate was centrifuged at 12000×*g* for 5 min at 4 ℃. His-LrhA protein was purified by Ni-NTA Resin (ThermoFisher, 88222) in a gravity-flow column, and the bound protein was eluted with PBS containing 250 mM imidazole. The eluted protein was analyzed by SDS-PAGE, and dialyzed at 4 °C in PBS with a 10 kDa dialysis tube (Millipore, UFC801096). Protein concentrations were determined by the Bradford assay (Beyotime, P0006C).

***Gene Ontology analysis***

For Gene Ontology analysis, the list of proteins was uploaded to the Gene Ontology database search portal (http://geneontology.org/) on molecular function to retrieve the plotted terms with their corresponding *P* values (Fisher's exact test with no correction). The raw *P*-value was determined by Fisher's exact test. The False Discovery Rate (FDR) was calculated by the Benjamini-Hochberg procedure. Only results for FDR and $P < 0.05$ were displayed.

***Motility test of E. coli lrhA$^+$ and lrhA$^-$ strains***

Tryptone swarm plates (1% tryptone, 0.5% NaCl and 0.3% Bacto agar) were spotted with 10 μL overnight culture of the indicated strains, and incubated for 4 h at 37 ℃, followed by overnight incubation at room temperature.

***Spacer acquisition assay***

*E. coli* strains carrying pT or pNT were grown overnight in LB media supplemented with ampicillin (100 μg mL$^{-1}$), then diluted to 1:50 times for further 36 h growth at 37 ℃ with shaking.

For experiments presented in Supplementary Fig. S10, strains were grown in fresh LB with ampicillin (100 μg mL$^{-1}$) (for pT plasmid maintenance) for 24 h and diluted 1:500 in fresh LB with ampicillin, grown for an additional 24 h. Passaging was performed for three days.

To monitor spacer acquisition, 200 μL of the cultures were collected by precipitation, washed thrice, and then resuspended in distilled water. These cells were used as the template for PCR amplification of the CRISPR array I by using the primer pair P71 (Adaptation-F) and P72 (Adaptation-R) as shown in Fig. 5d. All PCR products were run at 150 volts on a 2% agarose gels with GelStain (TransGen Biotech GS101) in Tris Acetate-EDTA buffer to identify parental and expanded arrays (parental array + 61bp). For sequencing newly acquired spacers, the expanded array bands were excised, and corresponding DNA fragments were purified using a PCR Clean-up kit (MN 740609.50), and ligated to T vectors (Takara, 6013).

***Electrophoretic mobility shift assay (EMSA)***

$P_{cas}$ (5 ng) probes were mixed with purified LrhA protein of different concentrations in the presence of binding buffer (10 mM Tris-HCl pH 7.5, 50 mM KCl, 1 mM DTT, 2.5% glycerol, 1 mM EDTA). A Ctrl probe generated by PCR from the *lacZα* gene with primers P69 and P70 was used as a control. The mixture was incubated at room temperature for 20 minutes, then separated by electrophoresis in 5% native polyacrylamide gels in 0.5×TBE buffer. Bands were transferred to PVDF membrane using the semi-dry transfer apparatus (Genscript) and visualized by Chemiluminescent Nucleic Acid Detection Module (ThermoFisher, 89880). Images were collected e d.

***Western blot***

To evaluate the expression of His-LrhA, 5 mL *E. coli* cells were grown to the indicated times, harvested by centrifugation and then resuspended in 0.5 mL lysis buffer (50 mM Tris-HCl, pH 7.5, 50 mM NaCl, 5% glycerol, 1 mM DTT, 20 μg $mL^{-1}$ lysozyme) containing the protease inhibitor cocktail (Roche, 04693132001). After sonication, the lysates were centrifuged at 13000×*g* for 15 min at 4 ℃. The cell extracts were added with SDS sample loading buffer and boiled for 5 min at 95 ℃, then 20 μL of the sample was loaded onto 12% Bio-Rad's stain-free SDS-PAGE gel (Bio-Rad, 1610185), and blotted onto PVDF membranes using the semi-dry transfer apparatus (Genscript). Total protein on the membrane was visualized by a stain-free blot module in a ChemDoc$^{TM}$ XRS$^{+}$ system. The membrane was blocked and incubated overnight with primary antibody (mouse anti-His Ab, Proteintech Group, Inc, 66005) at 4 ℃, followed by horseradish peroxidase (HRP)-conjugated secondary antibody for 1 h at room temperature. After washing, the immune complexes were detected by the ECL Western Blotting (Pierce, 32209). The image was visualized in the ChemDoc$^{TM}$ XRS+ system with Image Lab$^{TM}$ Software (Bio-Rad).

***Conjugation interference assays***

To assess type I-E CRISPR-Cas immunity by conjugation assay, a conjugative plasmid pTc, that carries a DNA fragment with four protospacer-adjacent motif (PAM)-containing spacers from pT [1], was used for evaluating CRISPR immunity during plasmid transfer. CRISPR-targeted conjugation plasmid (pTc) or CRISPR-non-targeted conjugation plasmid (pNTc) were transferred from *E. coli* WM3064 donor cells to wild-type (WT) or *lrhA* null mutants (Δ*lrhA*) of BW25113 and MC4100 strains (recipients) through conjugation. The frequency of plasmid conjugation was calculated as the ratio of transconjugants per total recipients. The relative CRISPR immunity was defined as the conjugation frequency of pNTc plasmid relative to that of pTc plasmid.

#### Fig. S1 | Sequence of the promoter of the *cas* operon ($P_{cas}$).

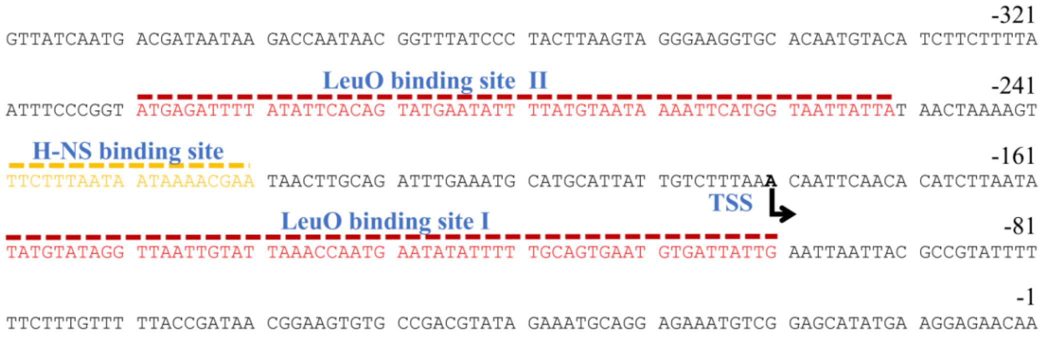


The sequence of *cas3*-*cse1* intergenic region (contains $P_{cas}$) is shown. Nucleotide positions are numbered relative to the first base of the *cse1* start codon. The H-NS and LeuO binding sites are indicated with yellow and red dashed lines respectively. TSS: Transcription Start Site. The source of the $P_{cas}$ sequence is from GenBank CP064677.1, *Escherichia coli* K-12 strain BW25113 chromosome (2877498 to 2877897).

**Fig. S2 | Filtering and enrichment of the $P_{cas}$-interacting proteomic data.**

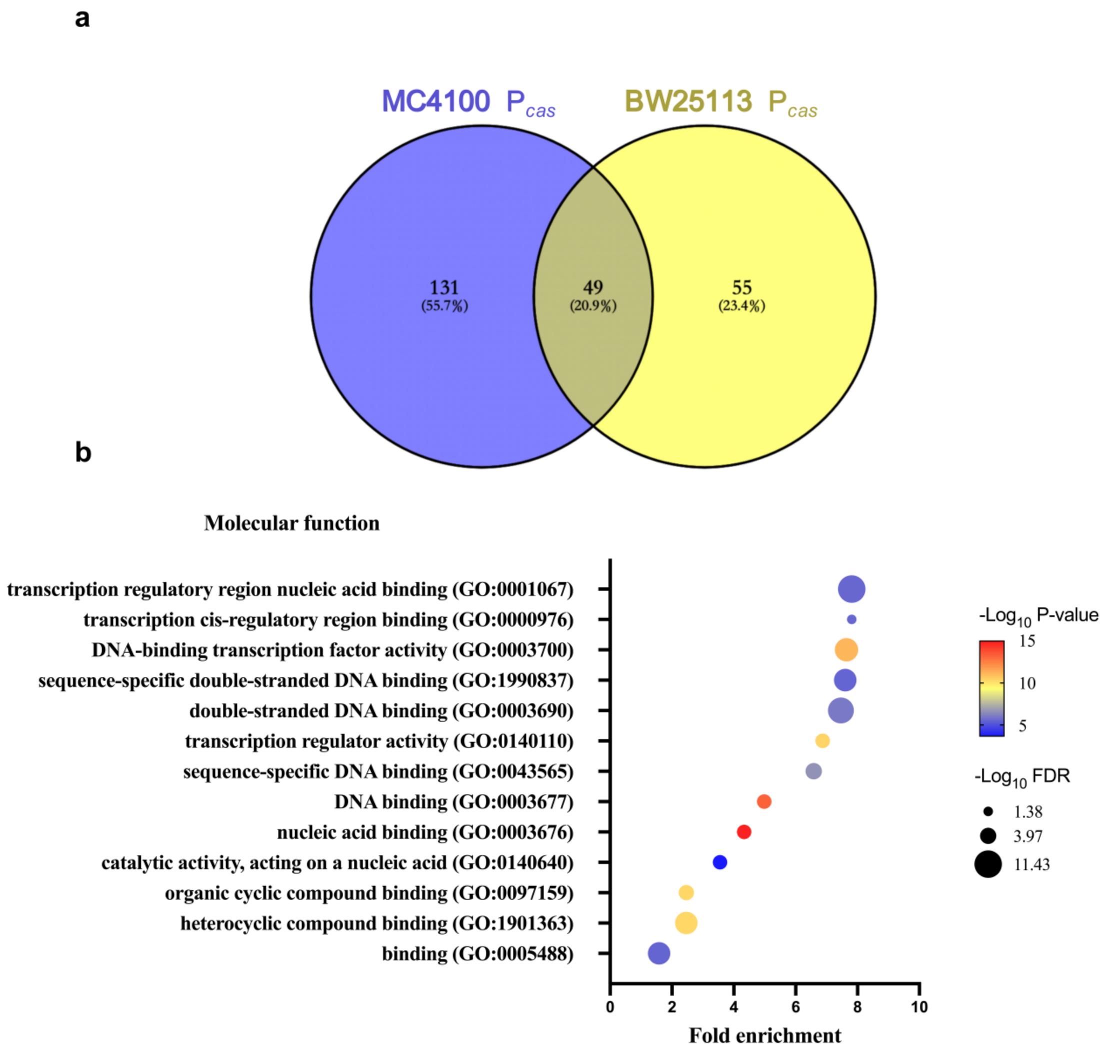


(**a**) After pull-down assay, mass spectrometry analysis of $P_{cas}$ binding proteins was performed as described in the main text. In *E. coli* MC4100 Δ*hns* mutant (ZJUTCBB0015), 442 proteins bound the biotinylated $P_{cas}$ probe ($P_{cas}$ group), and 359 proteins bound free biotin (Negative Control group, NC). In *E. coli* BW25113 Δ*hns* mutant (ZJUTCBB0009), 255 $P_{cas}$ group proteins and 228 NC proteins were detected**.** By removing proteins that appeared in the NC sample, we obtained 180 proteins (MC4100 $P_{cas}$) and 104 proteins (BW25113 $P_{cas}$) specifically bound to $P_{cas}$. Among them, 49 $P_{cas}$-binding proteins were shared by MC4100 $P_{cas}$ and BW25113 $P_{cas}$ group. (**b**) A bubble map was constructed based on Gene ontology (GO) to show subsets of $P_{cas}$-binding proteins in Supplementary Table S1. The significantly enriched 18 transcription factors (GO:0003700) are listed in Supplementary Table S2. Node color intensities with $-\log_{10}$ (FDR) and size scales with $-\log_{10}$ ($P$ value) were shown.

**Fig. S3 | Amino acid of peptides of LrhA sequenced by LC-MS/MS.**

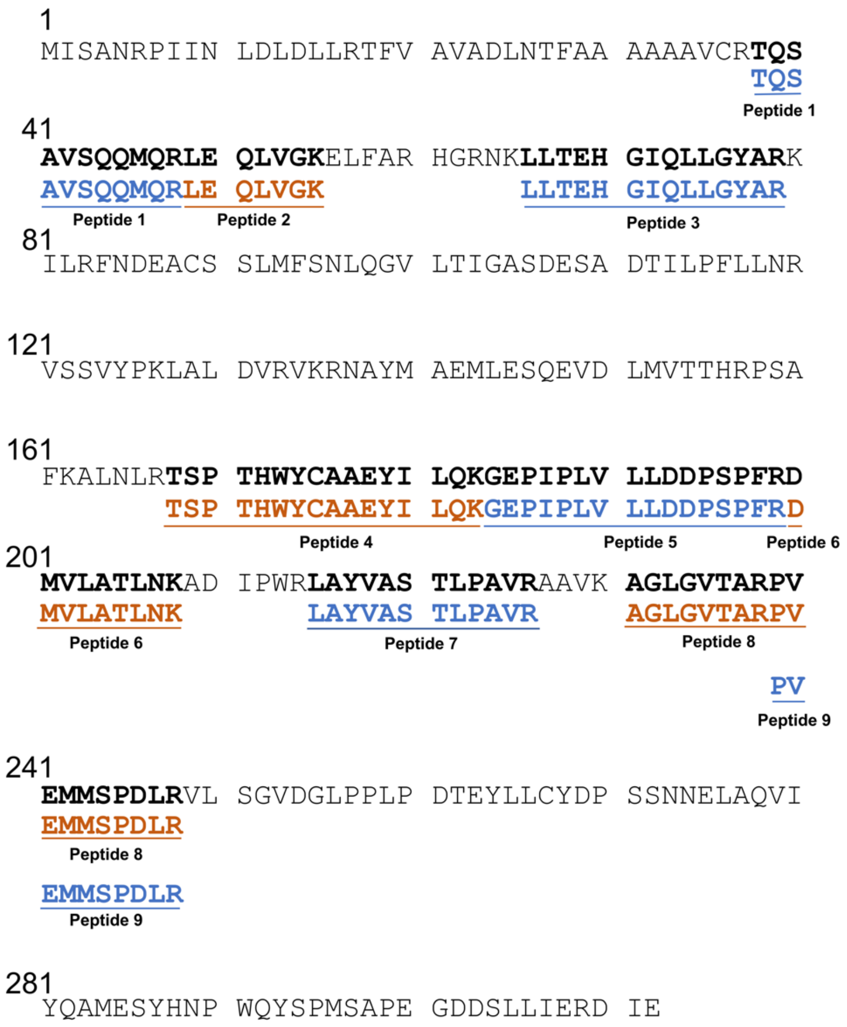


Peptide sequences identified by LC-MS/MS fragmentation spectrum of BW25113 Δ*hns* mutant sample were mapped to the amino acid sequence of LrhA.

**Fig. S4 | The effect of LrhA, LeuO, HexR, ArgP or SlyA on CRISPR immunity during conjugation.**

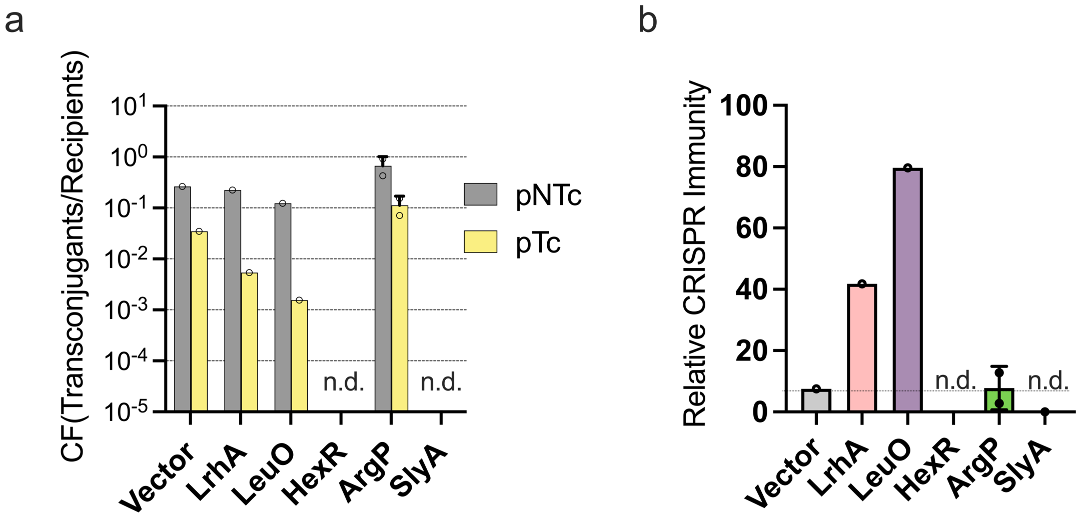


(**a**) Conjugation efficiency of CRISPR-targeted plasmid (pTc) and the CRISPR-non-targeted plasmid (pNTc) in *E. coli* BW25113 Δ*hns* Δ*leuO* (EC73) expressing LrhA, LeuO, HexR, ArgP or SlyA at 37 °C. (**b**) Type I-E CRISPR immunity in strains of (**a**). The relative CRISPR immunity was assessed by the conjugation frequency of pNTc plasmid relative to that of pTc plasmid. n.d.(not determined). Source data are provided as a Source Data file.

**Fig. S5 | Sequence of P$_{cas}$ in *E. coli* ER2738 strain (GenBank accession number, PP194457)**

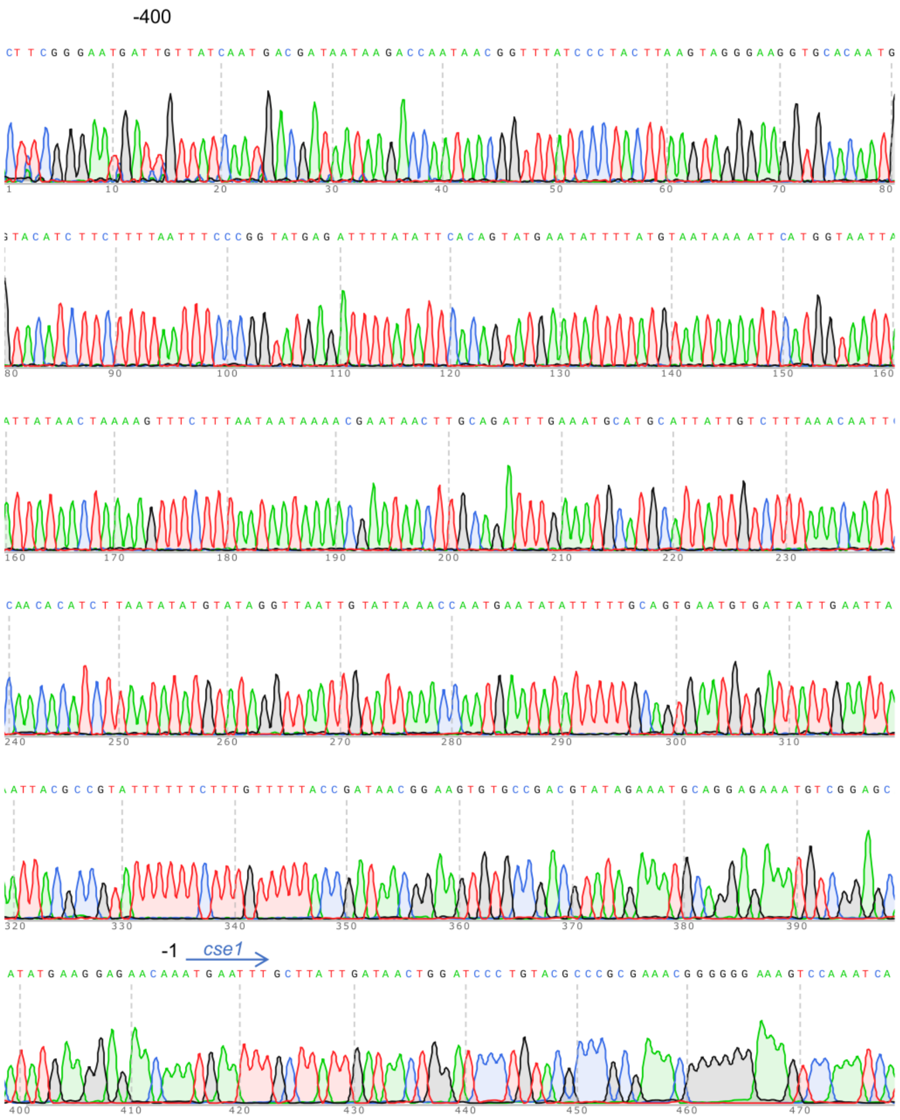


The sequence of P$_{cas}$ in *E. coli* K12 ER2738 strain was identical to that in typical *E. coli* K-12 strains.

**Fig. S6 | Motility of *E. coli lrhA*$^+$ and *lrhA*$^-$ strains.**

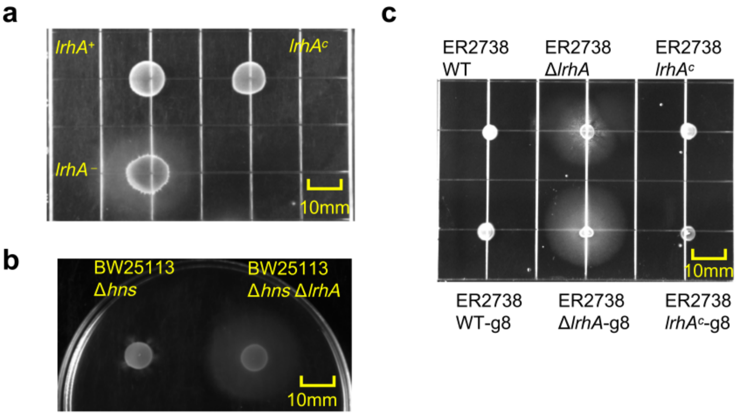


Motility assay in *E. coli lrhA*$^+$ and *lrhA*$^-$ strains. 10 µL overnight culture were spotted onto tryptone swarm agar (1% tryptone, 0.5% NaCl and 0.3% Bacto agar) plates, incubated for 4 h at 37°C and incubated at the room temperature (25°C) overnight. (**a**) Motility of BW25113 Δ*hns* Δ*leuO* (*lrhA*$^+$, EC73), Δ*hns* Δ*leuO* Δ*lrhA* (*lrhA*$^-$, EC77) or genomic *lrhA* complementation strain (*lrhA*$^c$, EC78). (**b**) Motility of BW25113 Δ*hns* (EC71) and Δ*hns* Δ*lrhA* (EC76) strains. (**c**) Motility of ER2738 wild-type (WT), Δ*lrhA* (EC87), genomic *lrhA* complementation strain (ER2738 *lrhA*$^c$, EC88) and their derivative strain with g8 spacer, WT-g8 (EC89), Δ*lrhA*-g8 (EC90), and *lrhA*$^c$-g8 (EC91).

**Fig. S7 | Effects of *lrhA* on the P*cas* activity in Δ*hns* Δ*leuO*, assessed by P*cas*-GFP reporter and qPCR assay.**

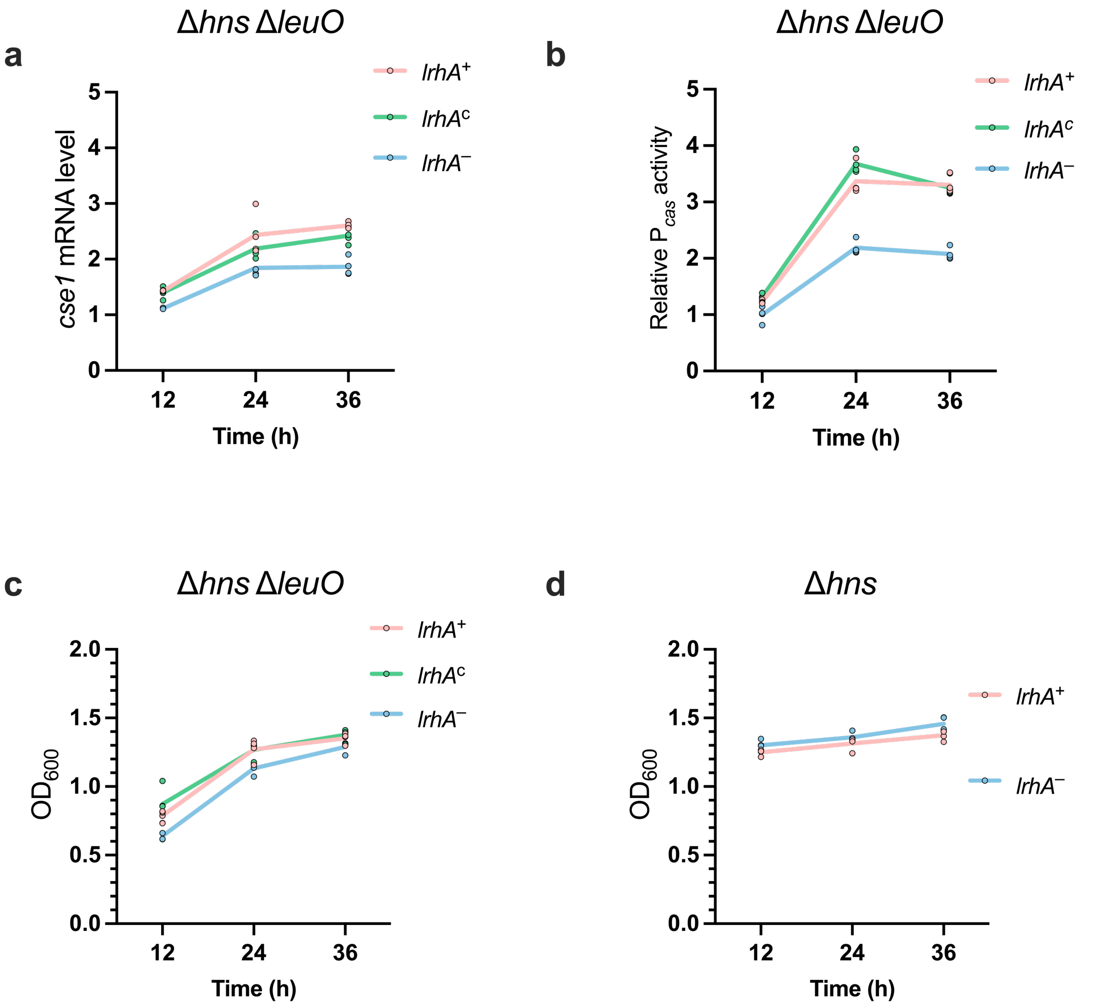


(**a**) Effects of genomic *lrhA* on the P*cas* activity assessed by qPCR assay in Δ*hns* Δ*leuO* at 37 °C. (**b**) Effects of genomic *lrhA* on the P*cas* activity assessed by P*cas*-*gfp* reporter assay in Δ*hns* Δ*leuO* at 37 °C. The P*cas* activity was indicated by GFP/$OD_{600}$, with the relative P*cas* activity of Δ*hns* Δ*leuO* Δ*lrhA* (*lrhA*$^{-}$, EC77) as 1 arbitrary unit. (**c**) Growth ($OD_{600}$) of the strain Δ*hns* Δ*leuO* (*lrhA*$^{+}$, EC73), Δ*hns* Δ*leuO* Δ*lrhA* (*lrhA*$^{-}$, EC77) and genomic *lrhA* complementation strain (*lrhA*$^{c}$, EC78) at 37 ℃. (**d**) Growth ($OD_{600}$) of the strain Δ*hns* (*lrhA*$^{+}$, EC71) and Δ*hns* Δ*lrhA* (*lrhA*$^{-}$, EC76) at 37 ℃. Individual biological replicates are shown ($n$ = 4).

**Fig. S8 | Inactivating *cas3* abolished CRISPR immunity against M13 phage.**

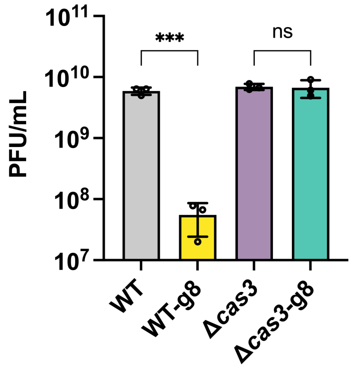


Quantification of plaque forming units (PFU) of M13 phage in ER2738 WT, ER2738 with g8 spacer (WT-g8, EC89), ER2738 Δ*cas3* (Δ*cas3*, EC96) ER2738 Δ*cas3* with g8 spacer (Δ*cas3*-g8, EC97) strains. Statistical significance was assessed using a One-way ANOVA. ***, $P < 0.001$. Bars are the means and error bars ± SD and individual biological replicates are shown ($n = 3$).

**Fig. S9 | LrhA does not affect CRISPR immunity against conjugation in *E. coli* BW25113 and MC4100 wild-type strains.**

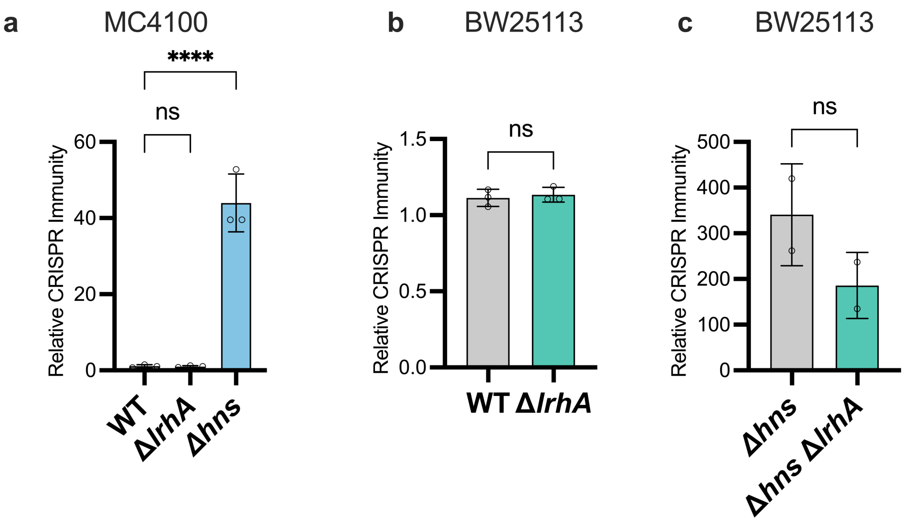


Type I-E CRISPR immunity assessed by conjugation assay in *E. coli* (**a**) MC4100 WT, Δ*lrhA* strain (EC92) and Δ*hns* strain (EC93), and in (**b**) BW25113 WT and Δ*lrhA* strain (EC75), and (**c**) BW25113 Δ*hns* (EC71) and Δ*hns* Δ*lrhA* (EC76) strain. The relative CRISPR immunity was defined as the conjugation frequency of pNTc plasmid relative to that of pTc plasmid. Statistical significance of the relative CRISPR immunity was assessed using One-way ANOVA for (**a**) or a two-tailed Student' s t-test for (**b** and **c**). ****, $P < 0.0001$. Bars are the means and error bars ± SD and individual biological replicates are shown ($n = 3$ for a and b, $n = 2$ for c).

**Fig. S10 | Spacer-acquisition assay.**

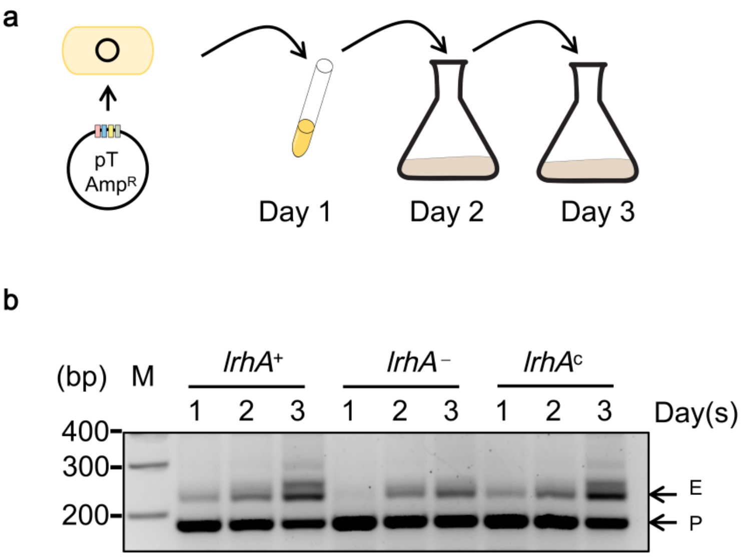


(**a**) Schematics of the spacer-acquisition assay. (**b**) Cultures of *E. coli* strains BW25113 Δ*hns* Δ*leuO* (*lrhA*$^{+}$, EC73), Δ*hns* Δ*leuO* Δ*lrhA* (*lrhA*$^{-}$, EC77) or genomic *lrhA* complementation strain (*lrhA*$^{c}$, EC78) harboring the pT plasmid were passaged for multiple days, and array expansion is assessed by PCR. Parental (P) and expanded (E) bands are indicated with arrows. (M: molecular weight marker). Gels are representative of two experiments yielding similar results.

**Fig. S11 | Effect of LrhA on CRISPR immunity assessed by plasmid loss in strains with *cas1-2* deletion or complementation.**

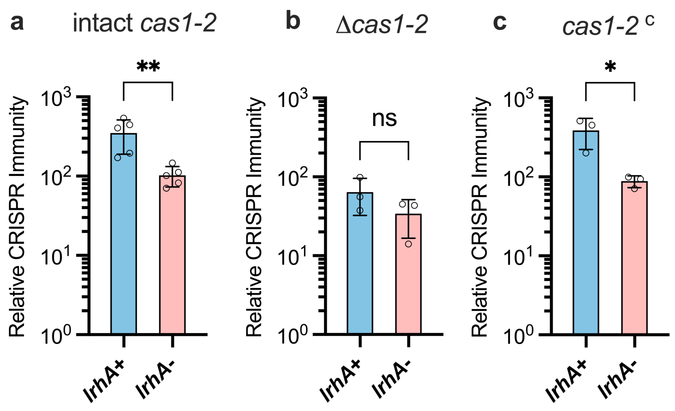


Plasmid loss assay of Δ*hns* Δ*leuO* (*lrhA*$^{+}$), Δ*hns* Δ*leuO* Δ*lrhA* (*lrhA*$^{-}$) with intact *cas1-2* (**a**) , *cas1-cas2* deletion (Δ*cas1-2*) (**b**) or its complementation (*cas1-2* $^{c}$) (**c**). Statistical significance was assessed using a two-tailed Student's t-test. *, *P* < 0.05 **, *P* < 0.01. Bars are the means and error bars denote standard deviation. Individual biological replicates are shown (*n* = 5 for **a**, *n* = 3 for **b** and **c**). *E. coli* strains BW25113 Δ*hns* Δ*leuO* (*lrhA*$^{+}$, EC73), Δ*hns* Δ*leuO* Δ*lrhA* (*lrhA*$^{-}$, EC77), Δ*hns* Δ*leuO* Δ*cas1-2* (*lrhA*$^{+}$ Δ*cas1-2*, EC81), Δ*hns* Δ*leuO* Δ*lrhA* Δ*cas1-2* (*lrhA*$^{-}$ Δ*cas1-2*, EC82), Δ*hns* Δ*leuO* *cas1-2* $^{c}$ (*lrhA*$^{+}$ *cas1-2* $^{c}$, EC84), Δ*hns* Δ*leuO*Δ*lrhA* *cas1-2* $^{c}$ (*lrhA*$^{-}$*cas1-2* $^{c}$, EC85), carrying pT or pNT were grown in antibiotic-free medium at 37 °C. After 36 h growth, plasmid stability of pT and pNT was evaluated by drop-plating on the LB plate supplemented with or without ampicillin. The relative CRISPR immunity was defined as the stability of the pNT plasmid relative to that of the pT plasmid.

**Fig. S12 | Schematic diagram of regulation of interference-adaptation feedback circuit by LrhA.**

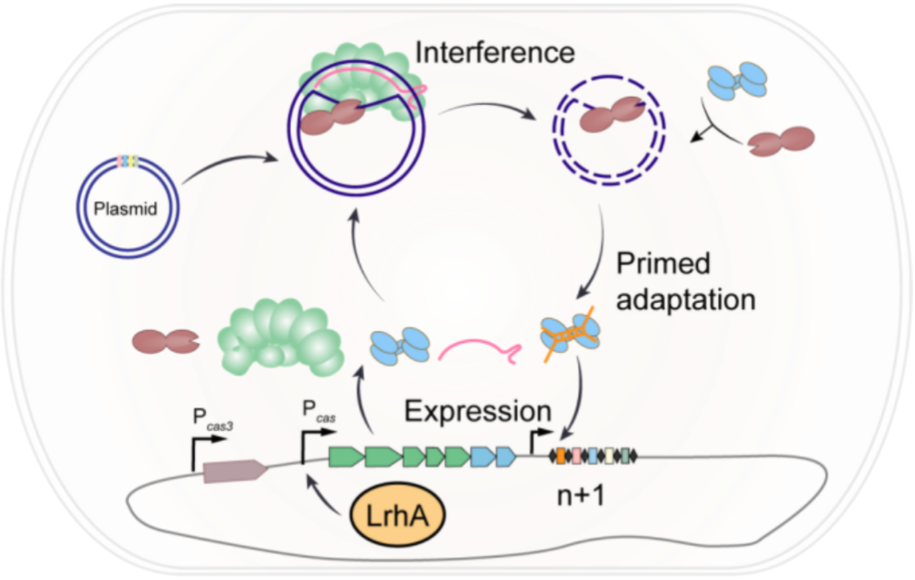


LrhA binds to $P_{cas}$ and activates transcription of the *cas* operon, increasing the amount of the surveillance complex Cascade and the integrase Cas1-Cas2. LrhA not only augments interference of CRISPR-targeted plasmid (pT), which is mediated by Cascade and the nuclease Cas3, but also promotes interference-driven spacer acquisition which is mediated by all *cas* proteins.

**Fig. S13 | Subfamily of LTTR proteins in *E. coli*.**

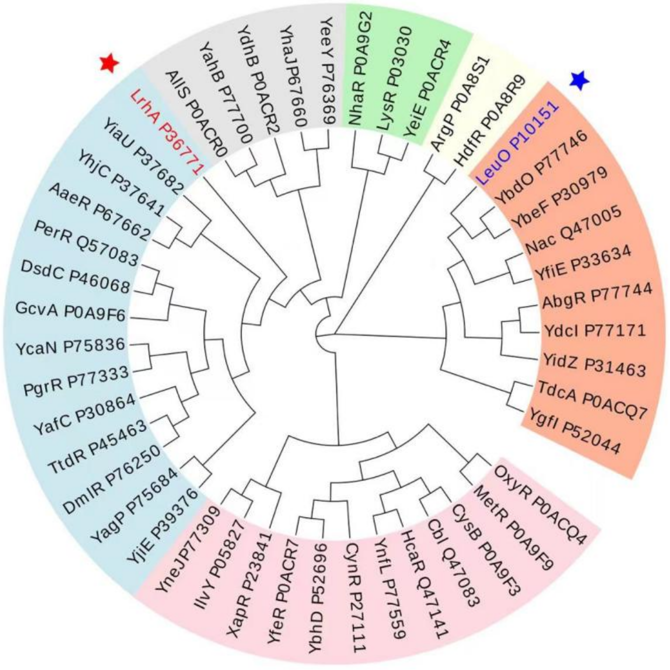


Phylogenetic analysis of 46 LTTRs from *E. coli*. Protein sequences were aligned with MEGA-X[2] software by using MUSCLE algorithms, and the phylogenetic tree was constructed by neighbour-joining method[3] and visualized by Evolview[4] tool. Bootstrap analysis was performed with 1,000 replicates to support each branch. All LTTRs were grouped into 6 clusters (subfamilies), as indicated by different background colors. A solid red star indicates LrhA, and a blue star indicates LeuO.

#### Fig. S14 | Phylogenetic tree of LeuO and LrhA protein.

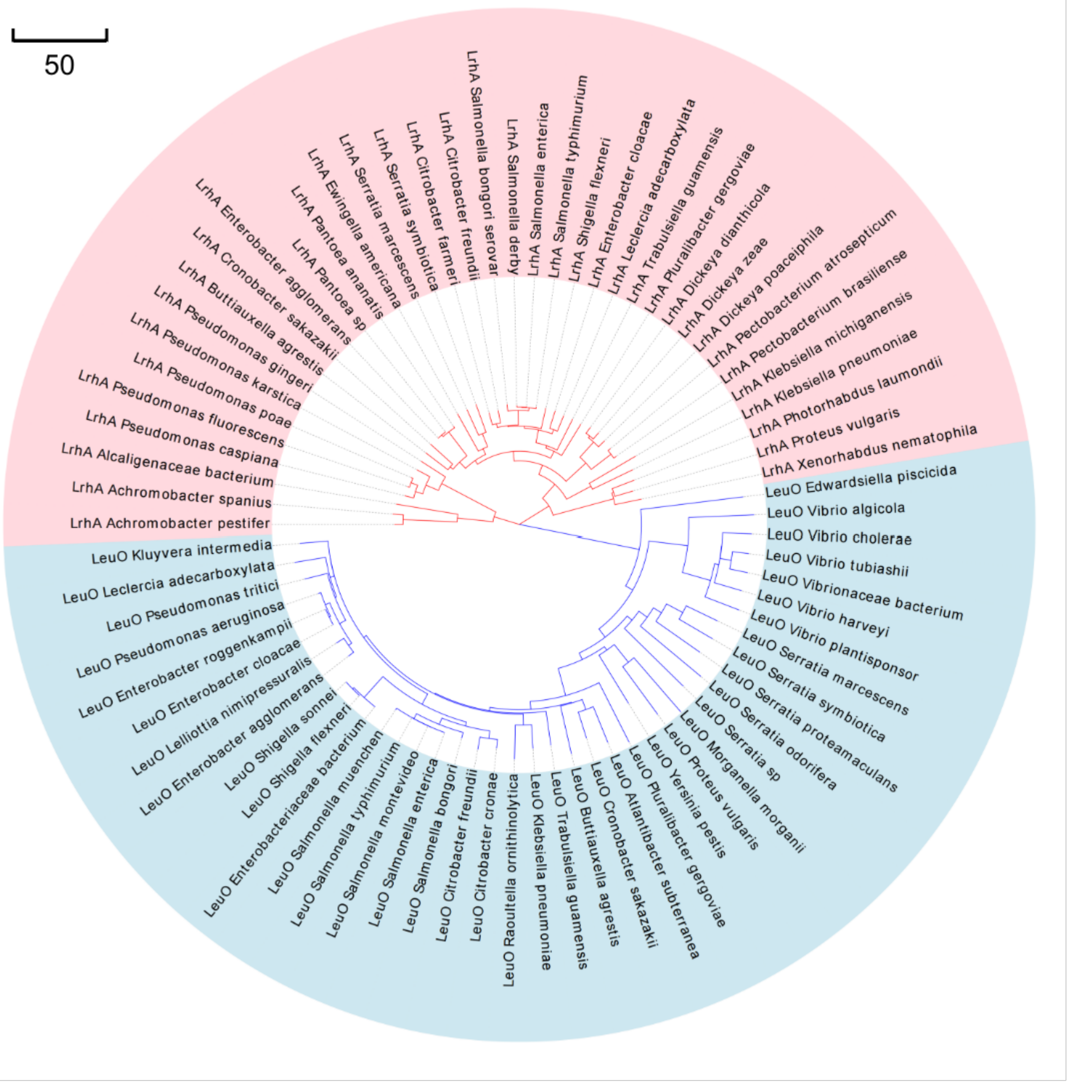


Phylogenetic tree of LeuO and LrhA protein sequences is built by using neighbour-joining method[3] with MEGA-X[2]. Bootstrap analysis was performed with 1, 000 replicates to support each branch[5]. LrhA proteins from 37 bacteria species are highlighted in pink, and LeuO proteins from 40 bacteria species are highlighted in blue. The optimal tree with the sum of branch length of 1484.43623324 is shown. The bar indicates the number of substitutions per site.

**Fig. S15 | Schematic illustration of pT (the CRISPR-targeted plasmid) and pNT (the CRISPR-non-targeted plasmid).**

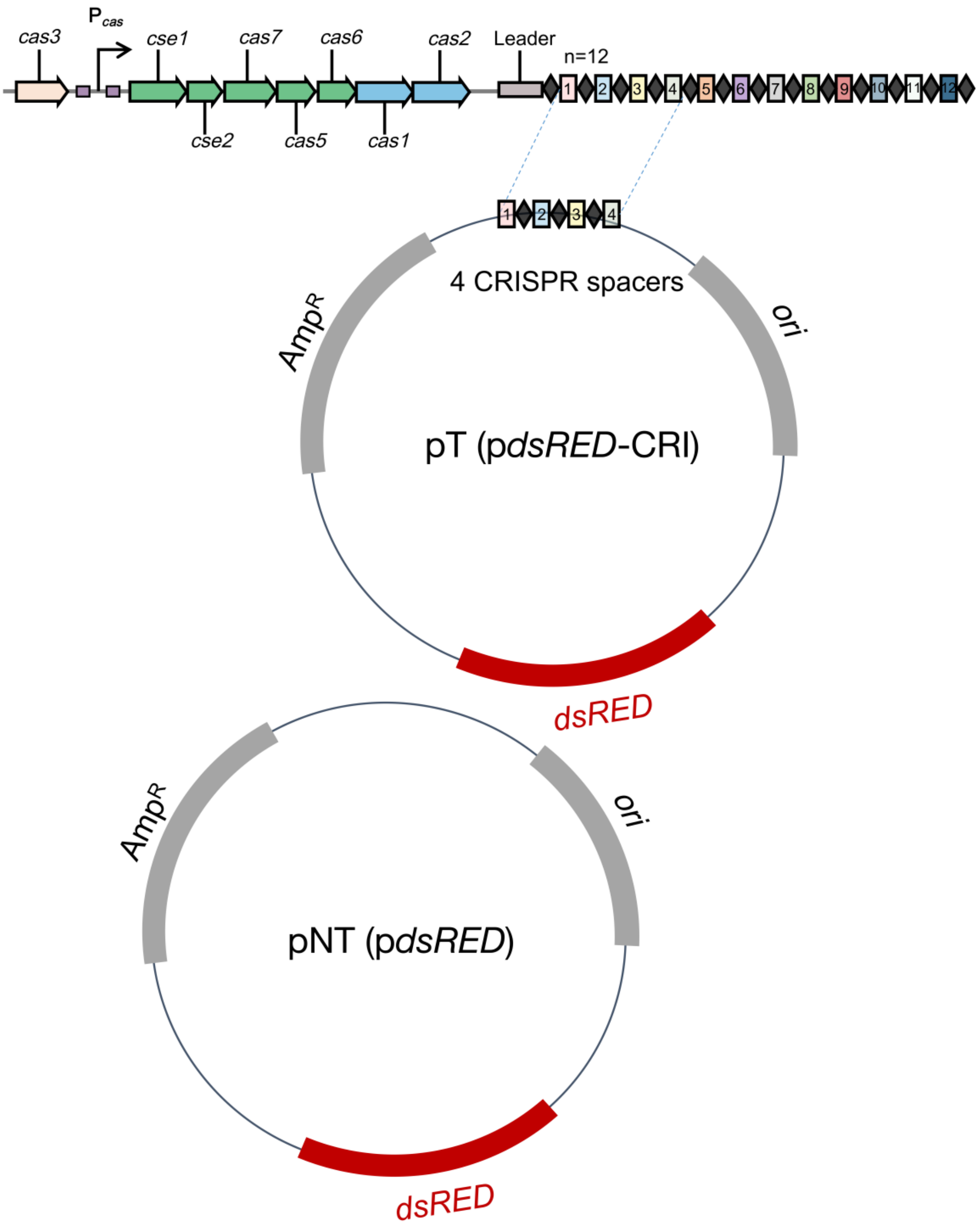


The 4 spacers in pT are from *E. coli* CRISPR array I locus, which can be targeted by the type I-E CRISPR-Cas system. The sequence of the 4 spacers is as follows. Repeats are indicated with letters on grey background, PAMs are purple letters and spacers 1-4 are indicated with red, blue, yellow and green letters.

AAGCTTTCGCAGACGCGCGGCGATACGCTCACGCAGAGTTCCCCGCGCCAGCGGGGATAAAAAGCAGCCGAAGCCAAAGGTGATGCCGAACACGCTGAGTTCCCCGCGCCAGCGGGGATAAAAAGGGCTCCCTGTCGGTTGTAATTGATAATGTTGAGAGTTCCCCGCGCCAGCGGGGATAAAAAGTTTGGATCGGGTCTGGAATTTCTGAGCGGTCGC

**Table S1 | GO Analysis of $P_{cas}$-binding transcription factors.**

| GO molecular function | Fold Enrichment | raw P-value | FDR[A] | Protein Number |
|---|---|---|---|---|
| transcription regulatory region nucleic acid binding (GO:0001067) | 7.81 | 2.53E-06 | 3.73E-12 | 9 |
| transcription cis-regulatory region binding (GO:0000976) | 7.81 | 2.53E-06 | 4.17E-02 | 9 |
| **DNA-binding transcription factor activity (GO:0003700)** | **7.64** | **7.49E-12** | **6.26E-09** | **18** |
| sequence-specific double-stranded DNA binding (GO:1990837) | 7.6 | 3.12E-06 | 2.06E-08 | 9 |
| double-stranded DNA binding (GO:0003690) | 7.46 | 9.56E-07 | 7.63E-11 | 10 |
| transcription regulator activity (GO:0140110) | 6.87 | 3.97E-11 | 6.51E-04 | 18 |
| sequence-specific DNA binding (GO:0043565) | 6.58 | 2.30E-07 | 8.22E-05 | 12 |
| DNA binding (GO:0003677) | 4.98 | 6.10E-14 | 6.34E-04 | 27 |
| nucleic acid binding (GO:0003676) | 4.33 | 1.49E-15 | 7.04E-04 | 32 |
| catalytic activity, acting on a nucleic acid (GO:0140640) | 3.55 | 2.16E-04 | 6.94E-04 | 11 |
| organic cyclic compound binding (GO:0097159) | 2.46 | 4.94E-11 | 2.99E-04 | 37 |
| heterocyclic compound binding (GO:1901363) | 2.46 | 4.94E-11 | 2.48E-08 | 37 |
| binding (GO:0005488) | 1.58 | 3.05E-06 | 2.47E-08 | 41 |

[A] FDR（False Discovery Rate) was calculated by the Benjamini-Hochberg procedure. Only results for FDR and *P* < 0.05 were displayed.

**Table S2 | $P_{cas}$-binding transcription factors identified by pull-down-based mass spectrometry.**

| Protein IDs | Description | Accession number | Mr (kDa) | Score (MC4100 Δhns) | No. of peptide matched (MC4100 Δhns) | Matched %Coverage (MC4100 Δhns) | Score (BW25113 Δhns) | No. of peptide matched (BW25113 Δhns) | Matched %Coverage (BW25113 Δhns) |
|---|---|---|---|---|---|---|---|---|---|
| **LrhA** | Probable HTH LysR -type transcriptional regulator LrhA | P36771 | 34.593 | 196.360 | 15 | 55.4 | 198.000 | 9 | 33.0 |
| StpA | DNA-binding protein StpA | P0ACG1 | 15.347 | 8.305 | 3 | 16.4 | 179.840 | 10 | 60.4 |
| YfhH | Uncharacterized HTH-type transcriptional regulator YfhH | P37767 | 30.707 | 76.803 | 8 | 35.8 | 88.813 | 8 | 33.7 |
| DeoR | Deoxyribose operon repressor | P0ACK5 | 28.548 | 59.434 | 8 | 29.4 | 69.396 | 7 | 25.4 |
| McbR | GntR family transcriptional regulator | P76114 | 47.358 | 207.200 | 12 | 67.4 | 58.256 | 5 | 31.2 |
| ArgP | HTH LysR -type transcriptional regulator ArgP | P0A8S1 | 33.471 | 106.350 | 12 | 44.4 | 47.794 | 6 | 22.9 |
| SlyA | Transcriptional regulator SlyA | P0A8W2 | 16.353 | 58.809 | 5 | 47.2 | 32.957 | 4 | 28.5 |
| OmpR | DNA-binding dual transcriptional regulator OmpR | P0AA16 | 27.353 | 70.659 | 14 | 56.5 | 31.332 | 4 | 18.0 |
| HexR | HTH rpiR-type transcriptional regulator HexR | P46118 | 31.957 | 61.697 | 8 | 36.3 | 28.843 | 3 | 13.8 |
| AllR | HTH iclR-type transcriptional repressor AllR | P0ACN4 | 29.269 | 28.704 | 9 | 32.5 | 26.560 | 4 | 14.8 |
| Rob | Right origin-binding protein | P0ACI0 | 33.144 | 43.399 | 9 | 34.6 | 25.026 | 3 | 10.7 |
| XylR | Xylose operon regulatory protein | P0ACI3 | 44.869 | 51.841 | 10 | 32.1 | 22.501 | 3 | 8.9 |
| AaeR | HTH LysR -type transcriptional activator AaeR | P67662 | 34.516 | 33.052 | 10 | 30.4 | 22.245 | 3 | 11.3 |
| KdgR | Transcriptional regulator KdgR | P76268 | 30.029 | 41.019 | 7 | 30.4 | 14.698 | 2 | 11.0 |
| YdcI | HTH LysR -type transcriptional regulator | P77171 | 33.373 | 34.342 | 4 | 24.1 | 13.989 | 2 | 7.2 |
| FnR | Fumarate and nitrate reduction regulatory protein | P0A9E5 | 27.967 | 5.503 | 2 | 13.6 | 11.937 | 2 | 11.2 |
| BluR | HTH merR-type transcriptional repressor BluR | P75989 | 28.218 | 7.029 | 2 | 10.3 | 11.619 | 2 | 11.1 |
| LexA | LexA repressor | P0A7C2 | 22.357 | 31.580 | 5 | 30.7 | 8.571 | 1 | 6.9 |

**Table S3 | Features of newly formed spacers**

| Bacterial strain | Clone number | DNA source | Spacer position | Spacer length | Position (complement strand)[B] | PAM[A] | Spacer sequence |
|---|---|---|---|---|---|---|---|
| | A1 | plasmid | S1 | 32 | (636-607) | AAG | GAGCCGTCCTGCAGGGAGGAGTCCTGGGTCAC |
| | A2 | plasmid | S1 | 32 | 112-143 | ATG | TGAGTTAGCTCACTCATTAGGCACCCCAGGCT |
| | A3 | plasmid | S1 | 32 | (2671-2640) | AAG | CGGTTAGCTCCTTCGGTCCTCCGATCGCTGTC |
| | A4 | plasmid | S1 | 32 | 922-953 | GTG | GAGCAGTACGAGCGCGCCGAGGGCCGCCACCA |
| BW25113 Δ*hns* Δ*leuO* pT | A5 | plasmid | S1 | 32 | (557-526) | AAG | CCCTCGGGGAAGGACAGCTTCTTGTAGTCGGG |
| | A6 | plasmid | S1 | 32 | (4017-3986) | AAG | TCAGAGGTGGCGAAACCCGACAGGACTATAAA |
| | A7 | plasmid | S1 | 32 | (2706-2675) | AAG | GCGAGTTACATGATCCCCCATGTTGTGCAAAA |
| | A8 | plasmid | S1 | 32 | (2283-2252) | AAG | GCAAAATGCCGCAAAAAAGGGAATAAGGGCGA |
| | A9 | plasmid | S1 | 32 | 1867-1898 | AAG | CGGATGCCGGGAGCAGACAAGCCCGTCAGGGC |
| | A10 | plasmid | S1 | 32 | 2007-2029 | TAA | AGGAGAAAATACCGCATCAGGCGCCATTCGCCA |
| | B1 | plasmid | S1 | 32 | (3841-3814) | AAG | ACACGATTTATCGCCACTTGCAGCAACCGCAG |
| | B2 | plasmid | S1 | 32 | (326-301) | AAG | CGCATGAACTCCTTGATGACGTCCTCGGAGGA |
| | B3 | plasmid | S1 | 32 | 3565-3596 | CAA | CCAGTTACCTTCGGAAAAAGAGTTGGTAACT |
| | B4 | plasmid | S1 | 32 | (4017-3986) | AAG | TCAGAGGTGGCGAAACCCGACAGGGCTATAAA |
| BW25113 Δ*hns* Δ*leuO* Δ*lrhA* pT | B5 | plasmid | S1 | 32 | (141-109) | AAG | CCTGGGGTGCCTAATGAGTGAGCTAACTCACA |
| | B6 | plasmid | S1 | 32 | (141-109) | AAG | CCTGGGGTGCCTAATGAGTGAGCTAACTCACA |
| | B7 | plasmid | S1 | 32 | (636-607) | AAG | GAGCCGTCCTGCAGGGAGGAGTCCTGGGTCAC |
| | B8 | plasmid | S1 | 32 | (2391-2360) | AAG | GATCTTACCGCTGTTGAGATCCAGTTCGATGT |
| | B9 | plasmid | S1 | 32 | (3960-3929) | AAG | CTCCCTCGTGCGCTCTCCTGTTCCGACCTTGC |
| | B10 | plasmid | S1 | 32 | (3960-3929) | AAG | CTCCCTCGTGCGCTCTCCTGTTCCGACCCTGC |

[A] Protospacer adjacent motif

[B] Based on plasmid pT sequence

**Table S4 | List of strains used in the study.**

| Strain | Description[A] | Source |
|---|---|---|
| BL21(DE3) | F- *ompT hsdS*$_B$ ($r_B^-$ $m_B^-$ ) *gal dcm*(DE3) | Tsingke Co., Ltd. |
| DH5α | $F^-$ø80(*lacZ*) ΔM15 Δ(*lacZYA-argF*) U169 *endA*1 *recA*1 *hsdR*17 ($r_k^-$, $m_k^+$ *sup*E44λ- *thi*-1 *gyrA*96 *relA*1 *phoA* | Tsingke Co., Ltd. |
| BW25113 | *lacI*$^q$ *rrnB*$_{T14}$ Δ *lacZ*$_{WJ16}$ *hsdR514* Δ*araBAD*$_{AH33}$ Δ*rhaBAD*$_{LD78}$ | [6] |
| MC4100 | F⁻ λ⁻ *araD139* Δ(*argF-lac*) *U169 rpsL150 relA deoC1 ptsF25 rbsR flbB5301* | [7] |
| ZJUTCBB0009 | BW25113 Δ*hns*$_{Km}$ | [8] |
| ZJUTCBB0015 | MC4100 Δ*hns*$_{Km}$ | [1] |
| EC71 | BW25113 Δ*hns*$_{FRT}$ | BW25113 Δ*hns*$_{Km}$ × pCP20 *flp* |
| EC72 | BW25113 Δ*leuO*$_{Cm}$ | Constructed by MY Fei |
| EC73 | BW25113 Δ*hns*$_{FRT}$ Δ*leuO*$_{FRT}$ | Constructed by MY Fei |
| EC74 | BW25113 Δ*cas*$_{Cm}$ | Constructed by YL Lu |
| EC75 | BW25113 Δ*lrhA*$_{FRT}$ | This study |
| EC76 | BW25113 Δ*hns*$_{FRT}$ Δ*lrhA*$_{FRT}$ | This study |
| EC77 | BW25113 Δ*hns*$_{FRT}$ Δ*leuO*$_{FRT}$ Δ*lrhA*$_{FRT}$ | This study |
| EC78 | EC77 FRT::*lrhA* | This study |
| EC79 | BW25113 Δ*hns*$_{FRT}$ Δ*leuO*$_{FRT}$ Δ*cas*$_{FRT}$ | This study |
| EC80 | BW25113 Δ*hns*$_{FRT}$ Δ*leuO*$_{FRT}$ Δ*cas*$_{FRT}$ Δ*lrhA*$_{FRT}$ | This study |
| EC81 | BW25113 Δ*hns*$_{FRT}$ Δ*leuO*$_{FRT}$ Δ(*cas1-cas2*)$_{FRT}$ | This study |
| EC82 | BW25113 Δ*hns*$_{FRT}$ Δ*leuO*$_{FRT}$ Δ*lrhA*$_{FRT}$ Δ(*cas1-cas2*)$_{FRT}$ | This study |
| EC83 | EC82 FRT::*lrhA* | This study |
| EC84 | EC81 FRT::(*cas1-cas2*) | This study |
| EC85 | EC82 FRT::(*cas1-cas2*) | This study |
| EC86 | EC83 FRT::(*cas1-cas2*) | This study |
| ER2738 | F′ *proA*$^+$*B*$^+$ *lacI*$^q$ Δ(*lacZ*)*M15 zzf::Tn10* (*Tet*$^R$)*/fhuA2 glnV thi* Δ(*lac-proAB*) Δ(*hsdMS-mcrB*)*5* (rk⁻ mk⁻ McrBC⁻ ) | New England Biolabs (E4104) |
| EC87 | ER2738 Δ*lrhA*$_{FRT}$ | This study |

| | | |
|---|---|---|
| EC88 | EC87 FRT::*lrhA* | This study |
| EC89 | ER2738 with g8 spacer (targeting phage M13 gene 8 encoding the coat protein) | This study |
| EC90 | EC87 with g8 spacer | This study |
| EC91 | EC88 with g8 spacer | This study |
| WM3064 | Donor strain for conjugation: *thrB*1004 *pro thi rpsL hsdS lacZ*ΔM15 RP4-1360 Δ(*araBAD*)567 Δ*dapA*1341::[*erm pir*(wt)] | [9] |
| EC92 | MC4100 Δ*lrhA*$_{FRT}$ | This study |
| EC93 | MC4100 Δ*hns*$_{Km}$ | This study |
| EC94 | ER2738 Δ*hns*$_{FRT}$ | This study |
| EC95 | ER2738 Δ*hns*$_{FRT}$ Δ*lrhA*$_{FRT}$ | This study |
| EC96 | ER2738 Δ*cas3*$_{FRT}$ | This study |
| EC97 | ER2738 Δ*cas3*$_{FRT}$ with g8 spacer | This study |

[A] Gene designations are Cm coding for chloramphenicol resistance, Km for kanamycin resistance. FRT is the Flp recombinase target site. Resistance cassettes flanked by FRT-sites were deleted using temperature-sensitive plasmid pCP20.

**Table S5 | List of plasmids used in the study.**

| Plasmids | Description | Reference or source |
|---|---|---|
| pGLO-$P_{cas}$-*gfp* | pGLO derivative, *gfp* expressed with $P_{cas}$, $Ap^R$ | [1] |
| pSU19 | p15A replicon, $Cm^R$ | [1] |
| pSU-$P_{BAD}$-*lrhA* | pSU19 derivative, *lrhA* expressed with $P_{BAD}$, $Cm^R$ | This study |
| pSU-$P_{BAD}$-*leuO* | pSU19 derivative, *leuO* expressed with $P_{BAD}$, $Cm^R$ | This study |
| pSU-$P_{BAD}$-*yfhH* | pSU19 derivative, *yfhH* expressed with $P_{BAD}$, $Cm^R$ | This study |
| pSU-$P_{BAD}$-*deoR* | pSU19 derivative, *deoR* expressed with $P_{BAD}$, $Cm^R$ | This study |
| pSU-$P_{BAD}$-*mcbR* | pSU19 derivative, *mcbR* expressed with $P_{BAD}$, $Cm^R$ | This study |
| pSU-$P_{BAD}$-*argP* | pSU19 derivative, *argP* expressed with $P_{BAD}$, $Cm^R$ | This study |
| pSU-$P_{BAD}$-*slyA* | pSU19 derivative, *slyA* expressed with $P_{BAD}$, $Cm^R$ | This study |
| pSU-$P_{BAD}$-*ompR* | pSU19 derivative, *ompR* expressed with $P_{BAD}$, $Cm^R$ | This study |
| pSU-$P_{BAD}$-*hexR* | pSU19 derivative, *hexR* expressed with $P_{BAD}$, $Cm^R$ | This study |
| pSU-$P_{BAD}$-*nadR* | pSU19 derivative, *nadR* expressed with $P_{BAD}$, $Cm^R$ | This study |
| pSU-$P_{BAD}$-*glpR* | pSU19 derivative, *glpR* expressed with $P_{BAD}$, $Cm^R$ | This study |
| pKD46 | Expressing Red recombinase, *repA101*(Ts) *oriR101*, $Ap^R$ | [6] |
| pCP20 | Expressing FLP recombinase, *repA101*(Ts) pSC101 *ori*, $Cm^R$, $Ap^R$ | [6] |
| pKD3 | FRT $Cm^R$ PS1 PS2 *ori*R6K, $Ap^R$ | [6] |
| pKD4 | FRT $Km^R$ PS1 PS2 *ori*R6K, $Ap^R$ | [6] |
| pET28a-His-*lrhA* | pET28a derivative, *lrhA* expressed by T7/*lac* promoter, $Km^R$ | This study |
| pGLO-$P_{cas}$ del LBSI-*gfp* | pGLO-$P_{cas}$-*gfp* with deleted *leuO* binding site I(LBSI) of $P_{cas}$ | This study |
| pGLO-$P_{cas}$ del LBSII-*gfp* | pGLO-$P_{cas}$-*gfp* with deleted *leuO* binding site II(LBSII) of $P_{cas}$ | This study |
| pGLO-$P_{cas}$ del LBSI&II-*gfp* | pGLO-$P_{cas}$-*gfp* with deleted *leuO* binding site I and II(LBSI and II) of $P_{cas}$ | This study |
| pNT (p*dsRED*) | Non-Targeted plasmid expressing red fluorescent protein, pUC19 derivative, pMB1 replicon, $Ap^R$ | [1] Fig. S15 |
| pT (p*dsRED*-CRI) | p*dsRED* derivative carrying 4 spacers targeted by the type I-E CRISPR-Cas system of *E. coli*, $Ap^R$ | [1] Fig. S15 |
| pHG101 | Conjugation plasmid, $Km^R$ | Genbank PP943005 |

| | | |
|---|---|---|
| pNTc | pHG101 derivative, non-targeted plasmid expressing red fluorescent protein, $Km^R$ | This study |
| pTc | pHG101 derivative, carrying 4 spacers targeted by the type I-E CRISPR-Cas system of *E. coli*, $Km^R$ | This study |

**Table S6 | List of oligonucleotides used in this study**

| Project/task | No. | Primer | Sequence (5'→ 3') | Description |
|---|---|---|---|---|
| Cloning [A] | P1 | Clone *lrhA* to pSU-F | GGGCTAGAAATAATTTTGTTTAACTTTAAGAAGGAGATATACAT**ATGATAAGTGCAAATCGTCCGATAA** | Constructing pSU-P$_{BAD}$-*lrhA* |
| | P2 | Clone *lrhA* to pSU-R | ACTCTAGAGGATCCCCGGGTACCGAGCTCGAATTC**TTACTCGATATCCCTTTCAATCAAC** | |
| | P3 | Clone *lrhA* to pET28a-F | GGCAGCAGCCATCATCATCATCATCACATG**ATGATAAGTGCAAATCGTCCGATAA** | Constructing of pET28a-His-*lrhA* |
| | P4 | Clone *lrhA* to pET28a-R | CTTTCGGGCTTTGTTAGCAGCCGGATCTTA**TTACTCGATATCCCTTTCAATCAAC** | |
| | P5 | Clone *yfhH* to pSU-F | TTAAGAAGGAGATATACAT**ATGAACGGCTTACTTCGTATCC** | Constructing pSU-P$_{BAD}$-*yfhH* |
| | P6 | Clone *yfhH* to pSU-R | GGTACCGAGCTCGAATTC**TCAGACCAGTTTTTCACCAGC** | |
| | P7 | Clone *deoR* to pSU-F | TTAAGAAGGAGATATACAT**ATGGAAACACGTCGCGAAGA** | Constructing pSU-P$_{BAD}$-*deoR* |
| | P8 | Clone *deoR* to pSU-R | GGTACCGAGCTCGAATTC**TTAATACATCAACTTAATGCGCTGCG** | |
| | P9 | Clone *mcbR* to pSU-F | TTAAGAAGGAGATATACAT**ATGCCTGGAACGGGAAAAATG** | Constructing pSU-P$_{BAD}$-*mcbR* |
| | P10 | Clone *mcbR* to pSU-R | GGTACCGAGCTCGAATTC**TTAACGATTGTATTGCTGGTATAAAATAGC** | |
| | P11 | Clone *argP* to pSU-F | TTAAGAAGGAGATATACAT**ATGAAACGCCCGGACTACAG** | Constructing pSU-P$_{BAD}$-*argP* |
| | P12 | Clone *argP* to pSU-R | GGTACCGAGCTCGAATTC**TTAATCCTGACGAAGGACTTTGTG** | |
| | P13 | Clone *slyA* to pSU-F | TTAAGAAGGAGATATACAT**ATGAAATTGGAATCGCCACTAG** | Constructing pSU-P$_{BAD}$-*slyA* |
| | P14 | Clone *slyA* to pSU-R | GGTACCGAGCTCGAATTC**TCACCCTTTGGCCTGTAACTC** | |
| | P15 | Clone *ompR* to pSU-F | TTAAGAAGGAGATATACAT**ATGCAAGAGAACTACAAGATTCTGG** | Constructing pSU-P$_{BAD}$-*ompR* |
| | P16 | Clone *ompR* to pSU-R | GGTACCGAGCTCGAATTC**TCATGCTTTAGAGCCGTCCG** | |
| | P17 | Clone *hexR* to pSU-F | TTAAGAAGGAGATATACAT**ATGAATATGCTGGAAAAAATCCAGT** | Constructing pSU-P$_{BAD}$-*hexR* |
| | P18 | Clone *hexR* to pSU-R | GGTACCGAGCTCGAATTC**TTAGCGATCGTCACTTAAATTAAGTAAC** | |
| | P19 | Clone *nadR* to pSU-F | TTAAGAAGGAGATATACAT**ATGTCGTCATTTGATTACCTG** | Constructing pSU-P$_{BAD}$-*nadR* |
| | P20 | Clone *nadR* to pSU-R | GGTACCGAGCTCGAATTC**TTATCTCTGCTCCCCCATCA** | |

| | | | | |
|---|---|---|---|---|
| | P21 | Clone *glpR* to pSU-F | TTAAGAAGGAGATATACAT**ATGAAACAAACACAACGTCACAAC** | Constructing pSU-$P_{BAD}$-*glpR* |
| | P22 | Clone *glpR* to pSU-R | GGTACCGAGCTCGAATTC**TCAGCACAGCTCCAGTTGAATAT** | |
| Inactivation of chromosomal Gene [B] | P23 | *CmFRT*-F | TGTAGGCTGGAGCTGCTTC | Amplifying donor DNA for generating Δ*lrhA* mutant by λ-RED recombineering method |
| | P24 | *CmFRT*-R | ATATGAATATCCTCCTTAGTTCC | |
| | P25 | Δ*lrhA*-H1F | CGTTTTGCTTCTTTCAGCAC | |
| | P26 | Δ*lrhA*-H1R | AAGCAGCTCCAGCCTACAATATTATCACTTACTGGCGGCTC | |
| | P27 | Δ*lrhA*-H2F | TAAGGAGGATATTCATATTCGACGGACGATAGATAATTCC | |
| | P28 | Δ*lrhA*-H2R | AAACCAGACCTGCCAGTAACAC | |
| | P29 | Δ*leuO*-H1F | TCGCCCGGCGAAGAGACG | Amplifying donor DNA from Δ*leuO::cat* |
| | P30 | Δ*leuO*-H2R | CTTTCAACTGCTGATGGCAG | |
| | P31 | Δ*cas*-F | GCTGGATAATCTGCCCGGTG | Amplifying donor DNA from Δ*cas::cat* |
| | P32 | Δ*cas*-R | TCGGTGGAAGCGACTAA | |
| | P33 | Complement *lrhA*-F | gccgccagtaagtgataatatatgATAAGTGCAAATCGTCC | Complement genomic *lrhA* |
| | P34 | Complement *lrhA*-R | GAAGCAGCTCCAGCCTACAttaCTCGATATCCCTTTCAATC | |
| | P35 | Δ(*cas1*-*cas2*)-H1F | GGGTTATTAGGGCTACTCGG | Amplifying donor DNA for generating Δ(*cas1*-*cas2*) mutant |
| | P36 | Δ(*cas1*-*cas2*)-H1R | AAGCAGCTCCAGCCTACATTTATTACACCTCAAtcaCAGTGG | |
| | P37 | Δ(*cas1*-*cas2*)-H2F | GGAACTAAGGAGGATATTCATATAAACAAAGAATTAGCTGATC | |
| | P38 | Δ(*cas1*-*cas2*)-H2R | TCTTGGGAATAAGGATGG | |
| | P39 | Δ*cas3*-H1F | ATATGTACTTTCTTCCAGCTTTGG | Amplifying donor DNA for generating Δ*cas3* mutant |
| | P40 | Δ*cas3*-H1R | AAGCAGCTCCAGCCTACATAATAGCCTCCCTGTTTTTTAG | |
| | P41 | Δ*cas3*-H2F | GGAACTAAGGAGGATATTCATATCTTCGGGAATGATTGTTATC | |
| | P42 | Δ*cas3*-H2R | ACACCGTTGAACGAAGATCG | |
| Genotyping | P43 | Δ*lrhA*-test-F | GGGAACCAGTCAGAGGA | Confirming Δ*lrhA* constructs |
| | P44 | Δ*lrhA*-test-R | GAGCTTGCCCATAAACA | |
| | P45 | Δ*hns*-1500F | GCATTGCCCTTCTGGGGCCG | |

| | | | | |
|---|---|---|---|---|
| | P46 | Δ*hns*-999R | AGGAACCAGGATGTTGCCGG | Confirming Δ*hns* constructs |
| | P47 | Δ*leuO*-test-F | GATGTGCATTGGCGAAGT | Confirming Δ*leuO* constructs |
| | P48 | Δ*leuO*-test-R | GTCACGTAAGCGTCTCCCT | |
| | P49 | Δ*cas*-test-F | TCGCTTGATGAACTGATAGG | Confirming Δ(*casA*-*cas2*) constructs |
| | P50 | Δ*cas*-test-R | TCTTGGGAATAAGGATGG | |
| | P51 | Δ(*cas1*-*cas2*)-test-F | GGGTTATTAGGGCTACTCGG | Confirming Δ(*cas1*-*cas2*) constructs |
| | P52 | Δ(*cas1*-*cas2*)-test-R | TCGGTGGAAGCGACTAA | |
| | P53 | Δ*cas3*-test-F | GCTGGATAATCTGCCCGGTG | Confirming Δ*cas3* constructs |
| | P54 | Δ*cas3*-test-R | TATTGTTGCCACCCCTGATT | |
| Mutagenesis[C] | P55 | Delete $P_{cas}$ LBSI-F | **aattcaacacatcttaata**tgaattaattacgccgtatttttctttg | Constructing pGLO-$P_{cas}$ del LBSI-*gfp* and pGLO-$P_{cas}$ del LBSI&II-*gfp* |
| | P56 | Delete $P_{cas}$ LBSI-R | atattaagatgtgttgaattgtttaaagac | |
| | P57 | Delete $P_{cas}$ LBSII-F | **tcttcttttaatttcccggt**taactaaaagtttctttaataataaaacgaataacttgc | Constructing pGLO-$P_{cas}$ del LBSII-*gfp* and pGLO-$P_{cas}$ del LBSI&II-*gfp* |
| | P58 | Delete $P_{cas}$ LBSII-R | **attaaagaaacttttagtta**accgggaaattaaaagaagatgtacattg | |
| EMSA Probes | P67 | $P_{cas}$-F | TCTTCTTTTAATTTCCCGGT | Generating $P_{cas}$ probes for EMSA [D] |
| | P68 | $P_{cas}$-R | ATTCAATAATCACATTCACTGC | |
| | P69 | Ctrl Probe-F | GTCGTGACTGGGAAAACCCTGGCG | Generating Ctrl probe by PCR from *lacZα* gene |
| | P70 | Ctrl Probe-R | CCGCACAGATGCGTAAGGAG | |
| Checking expansion of CRISPR I array | P71 | Adaptation-F | AAATGTTACATTAAGGTTGGTG | Forward primer on CRISPR1 leader |
| | P72 | Adaptation-R | CATCACCTTTGGCTTCGGCTG | Reverse primer on CRISPR1 spacer 2 |
| qPCR | P73 | qPCR-*cse1*-F | GCCCCGTGACGATATGGA | |

| | | |
|---|---|---|
| P74 | qPCR-*cse1*-R | CGATAATTTGCCCAATGCAA |
| P75 | qPCR-*rpoD*-F | GACGAAGAAGATGGCGATGACGAC |
| P76 | qPCR-*rpoD*-R | TTCCTGAGCGGTAGCGTGACTG |

**[A] Target-gene specific sequences are in bold letters.**

**[B] Annealing region in primers for assembly of DNA sequences by overlap-extension PCR**

**[C] Position of mutations/deletions are marked in bold letters.**

**[D] Generating EMSA probes $P_{cas}$ FL, $P_{cas}$del LBSI, $P_{cas}$ del LBSII, $P_{cas}$ short probe by PCR from pGLO-$P_{cas}$-*gfp,* pGLO-$P_{cas}$ del LBSI-*gfp,* pGLO-$P_{cas}$ del LBSII-*gfp* and pGLO-$P_{cas}$ del LBSI&II-*gfp***
